## Supplementary figures and images for "Metapangenomics of the oral microbiome provides insights into habitat adaptation and cultivar diversity"

### Additional File 1

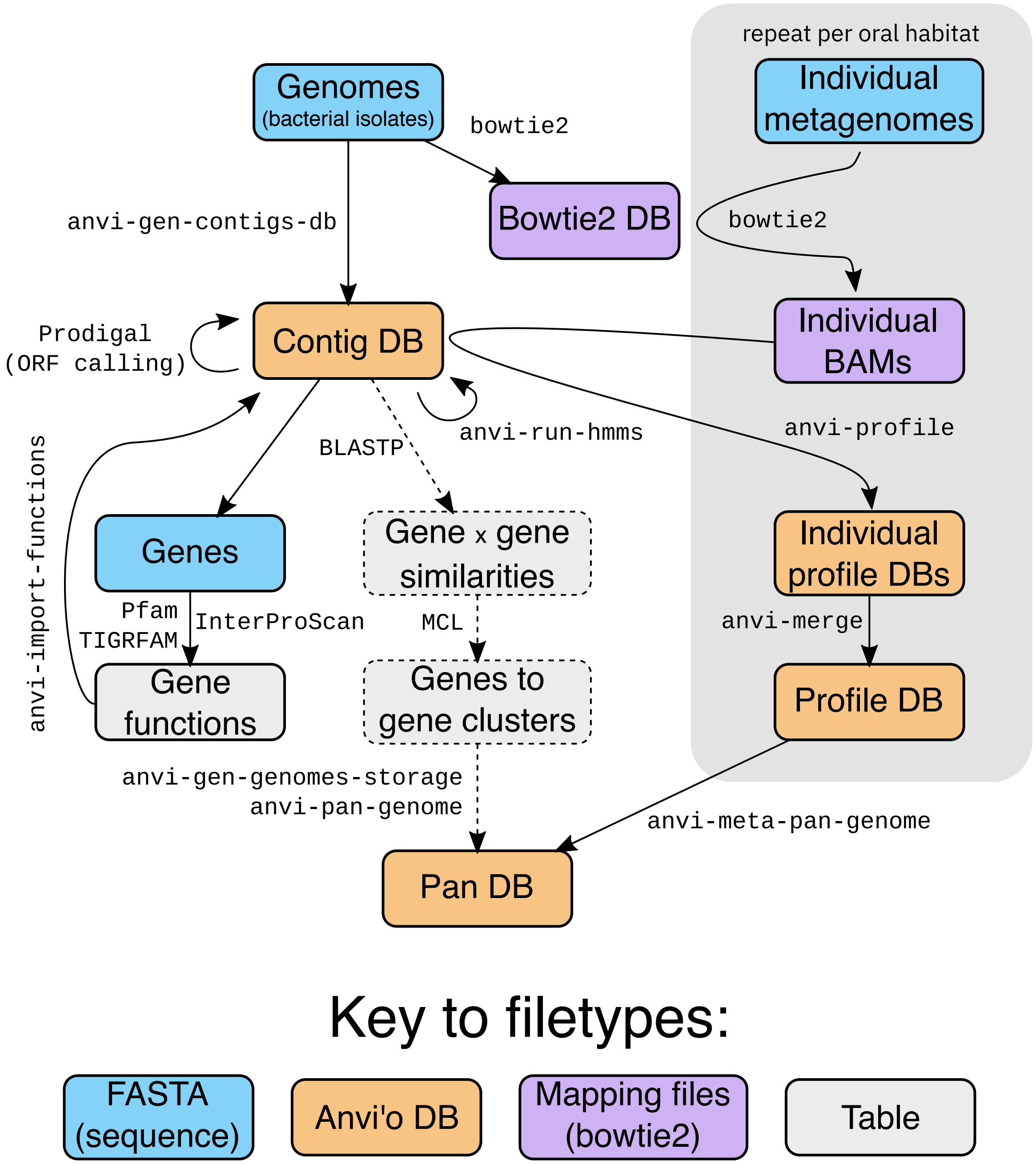

### Additional File 2

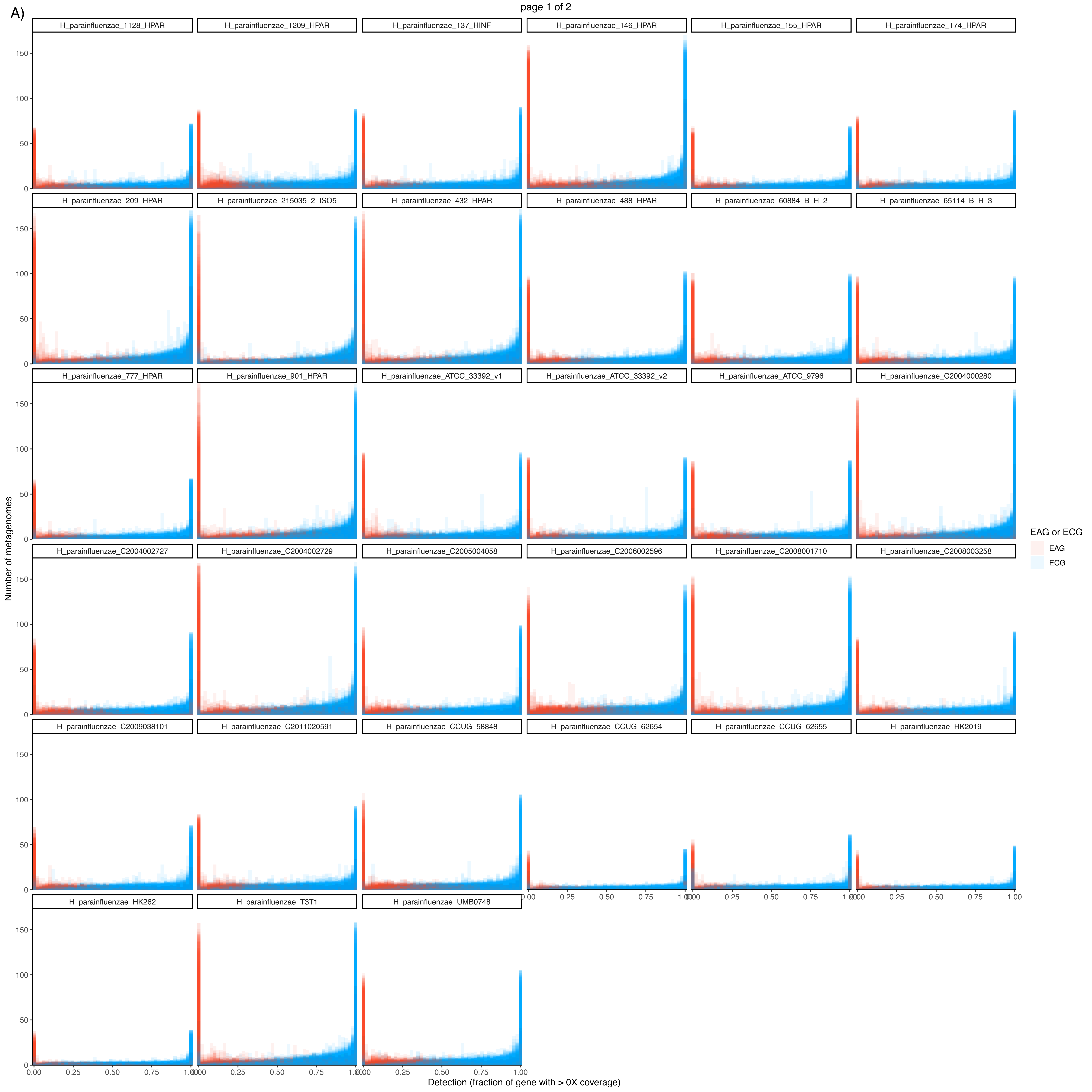

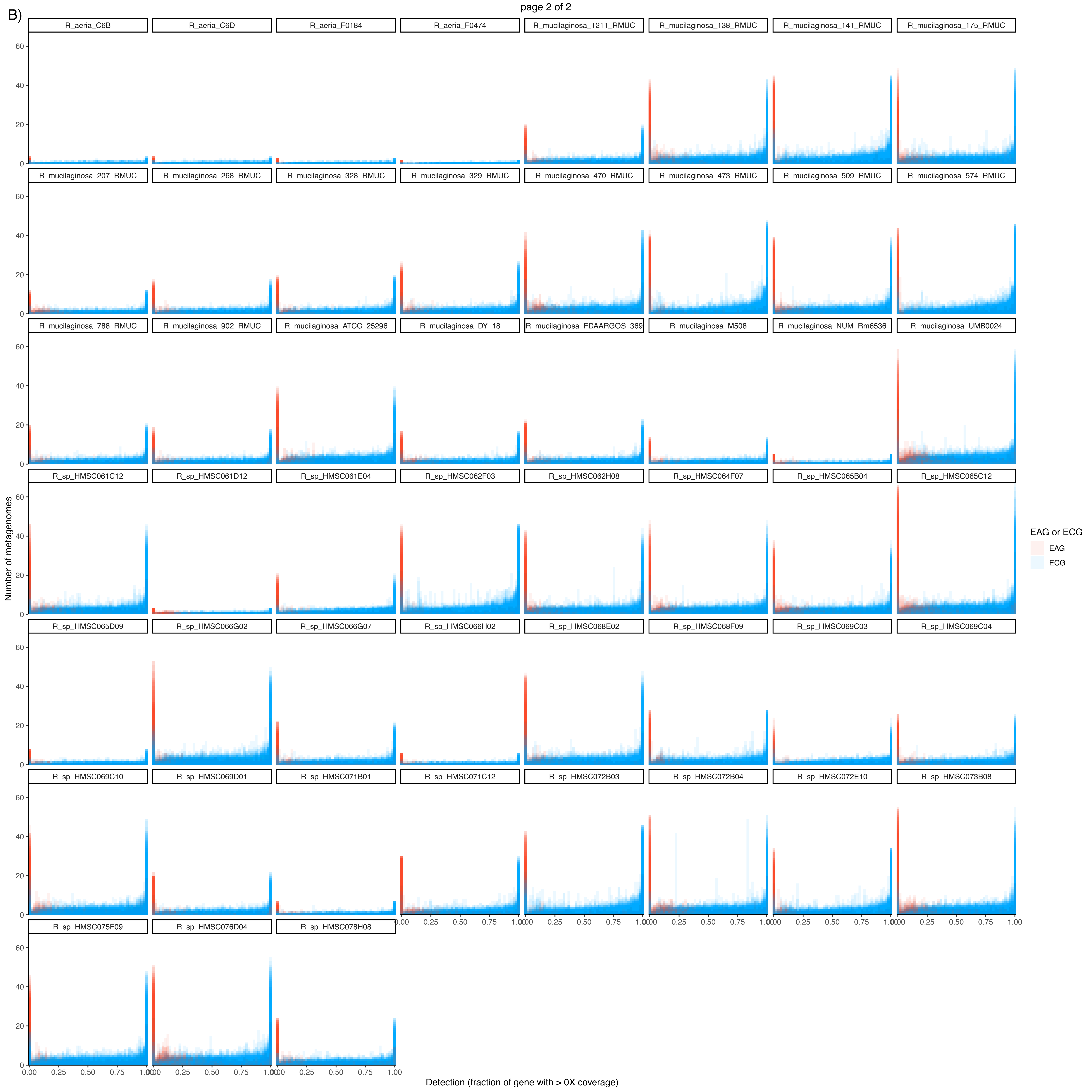

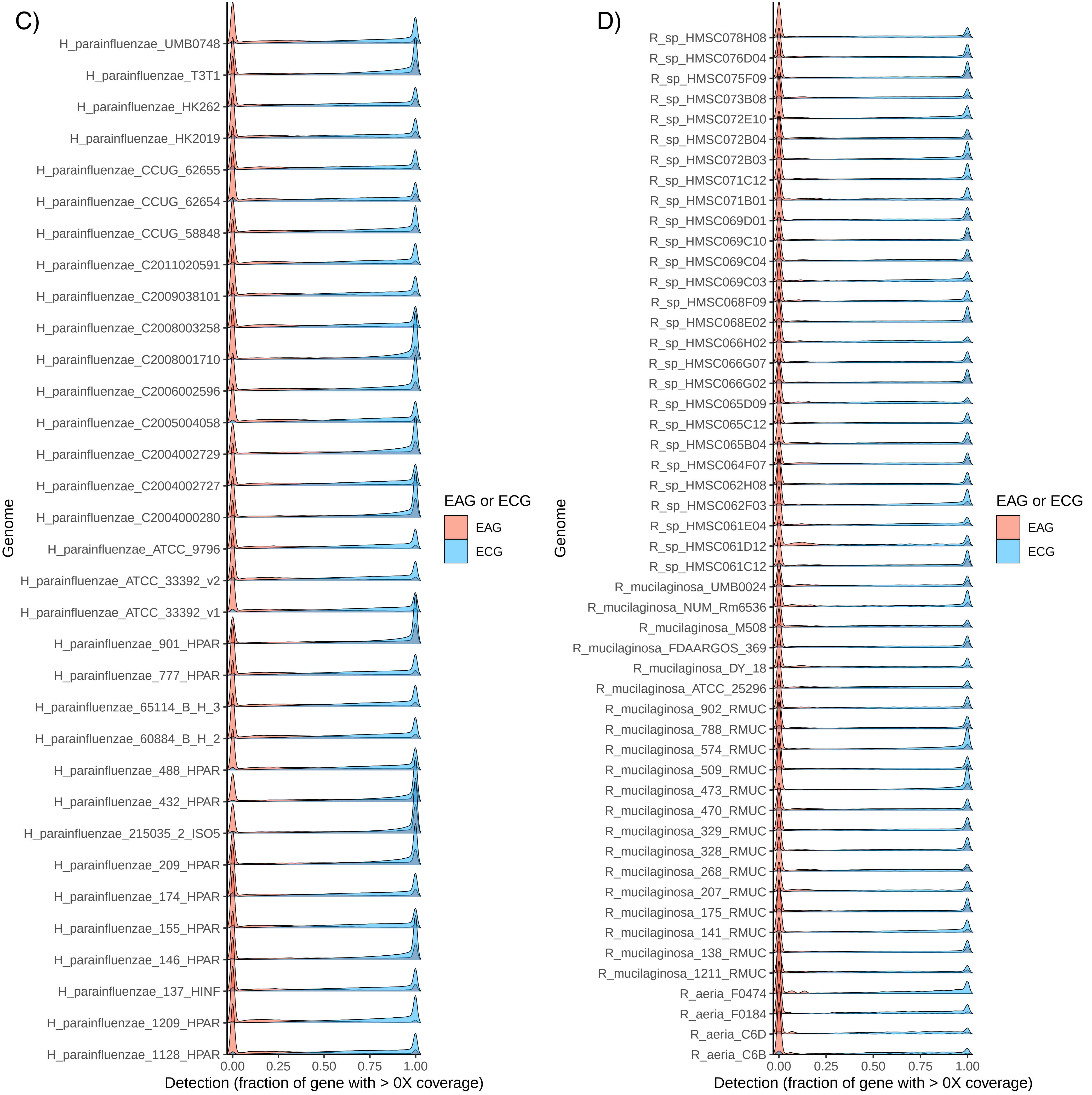

### Additional File 5

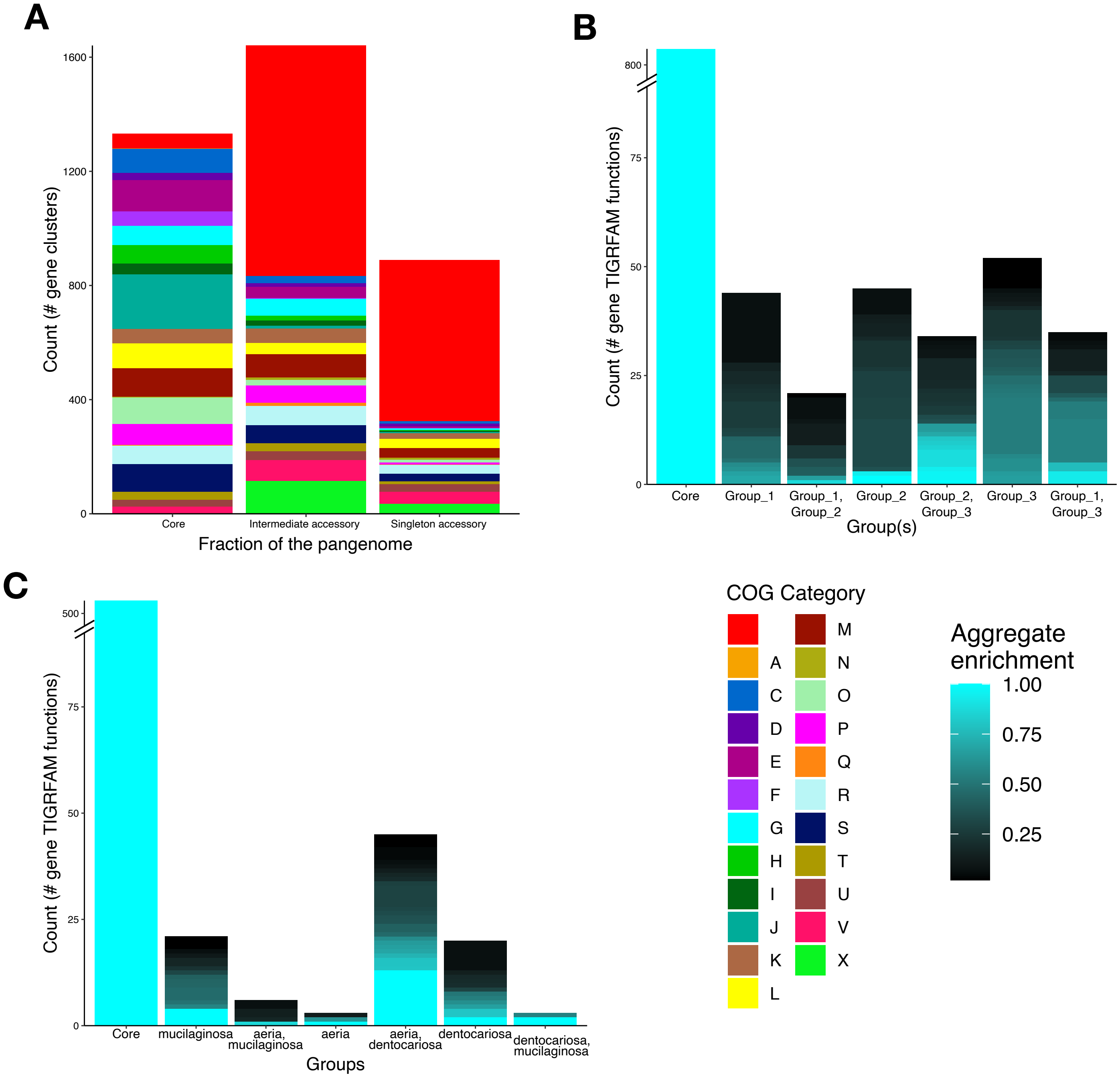

### Additional File 6

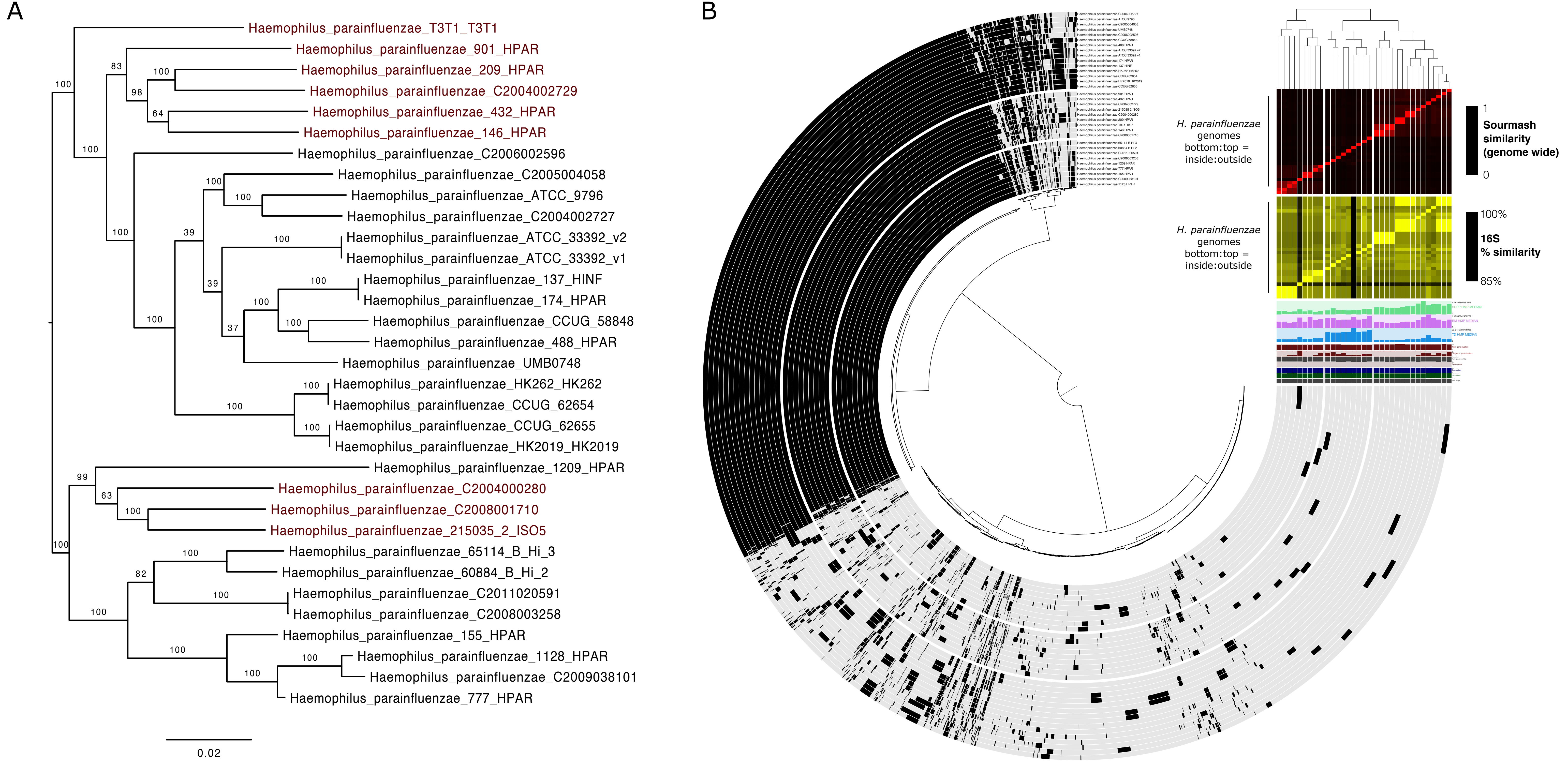

### Additional File 9

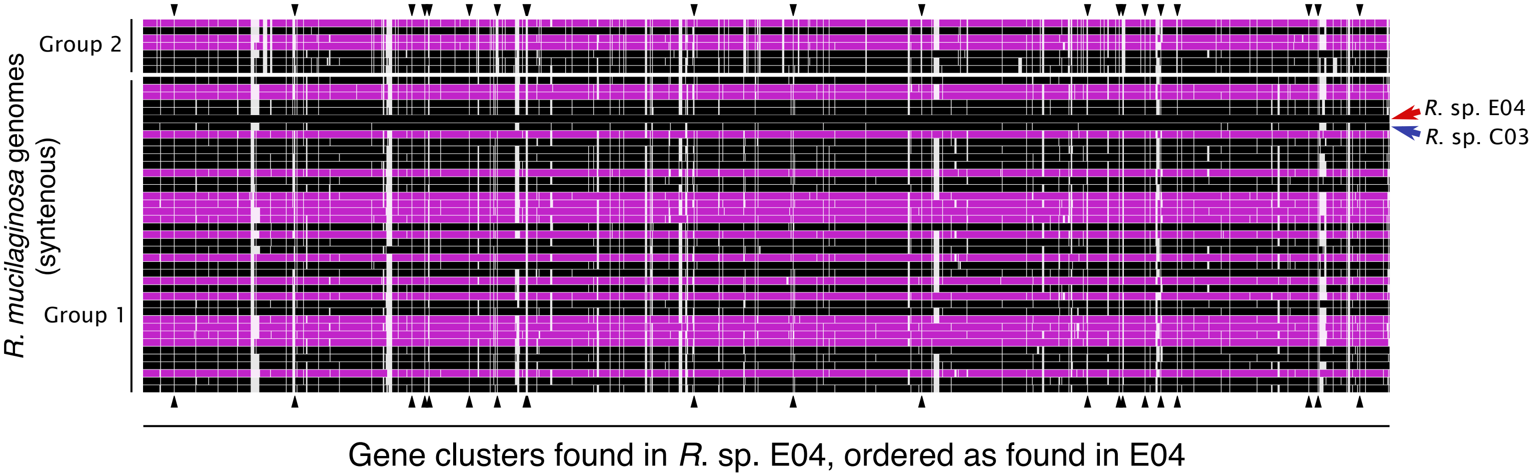

### Additional File 11

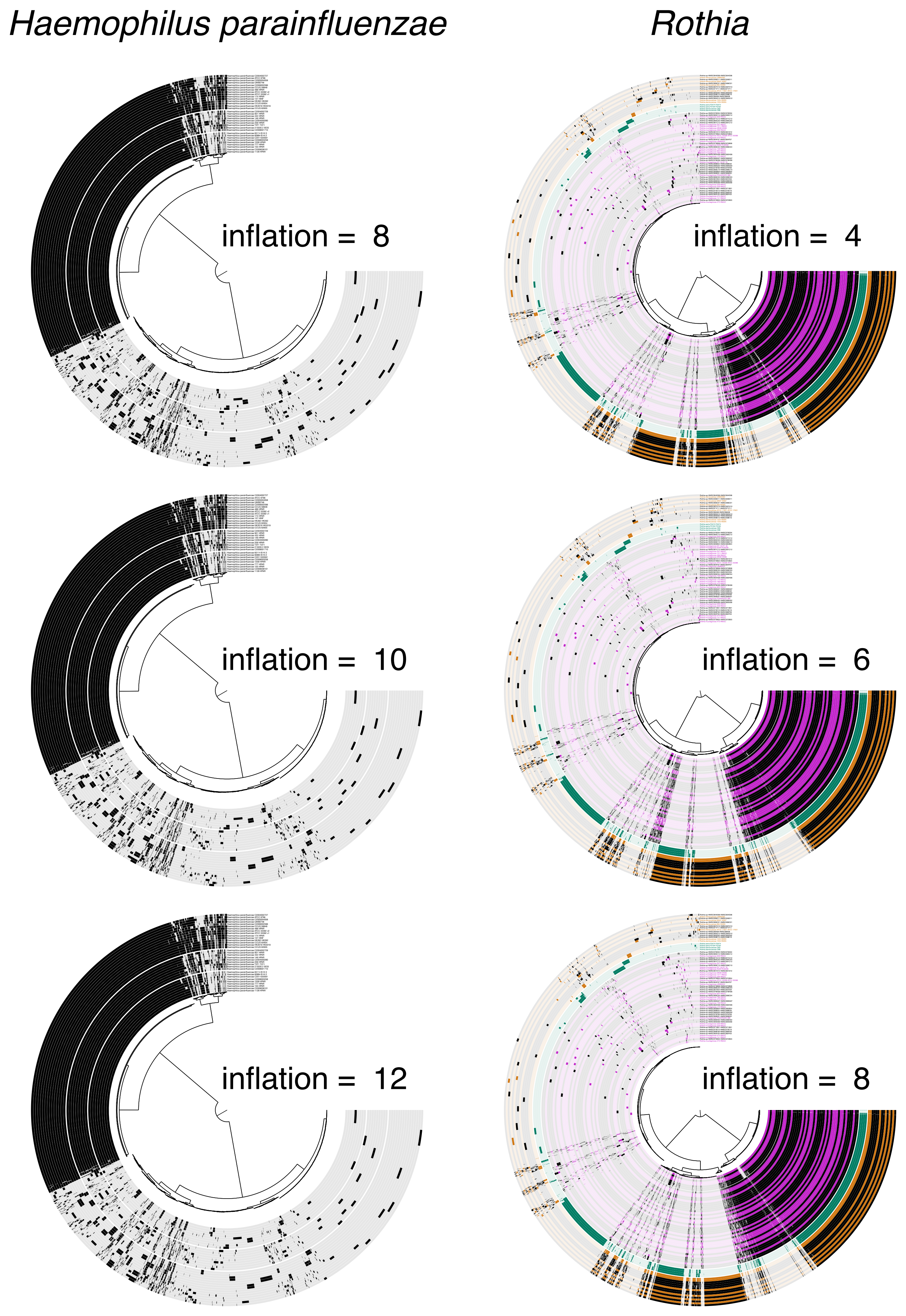
