## Additional File 10 for "Metapangenomics of the oral microbiome provides insights into habitat adaptation and cultivar diversity"

### Rothia_sp_HMSC069C03_HMSC069C03

Items order: <> User order (D: none; L: none) | Current view: single | Samples order: custom

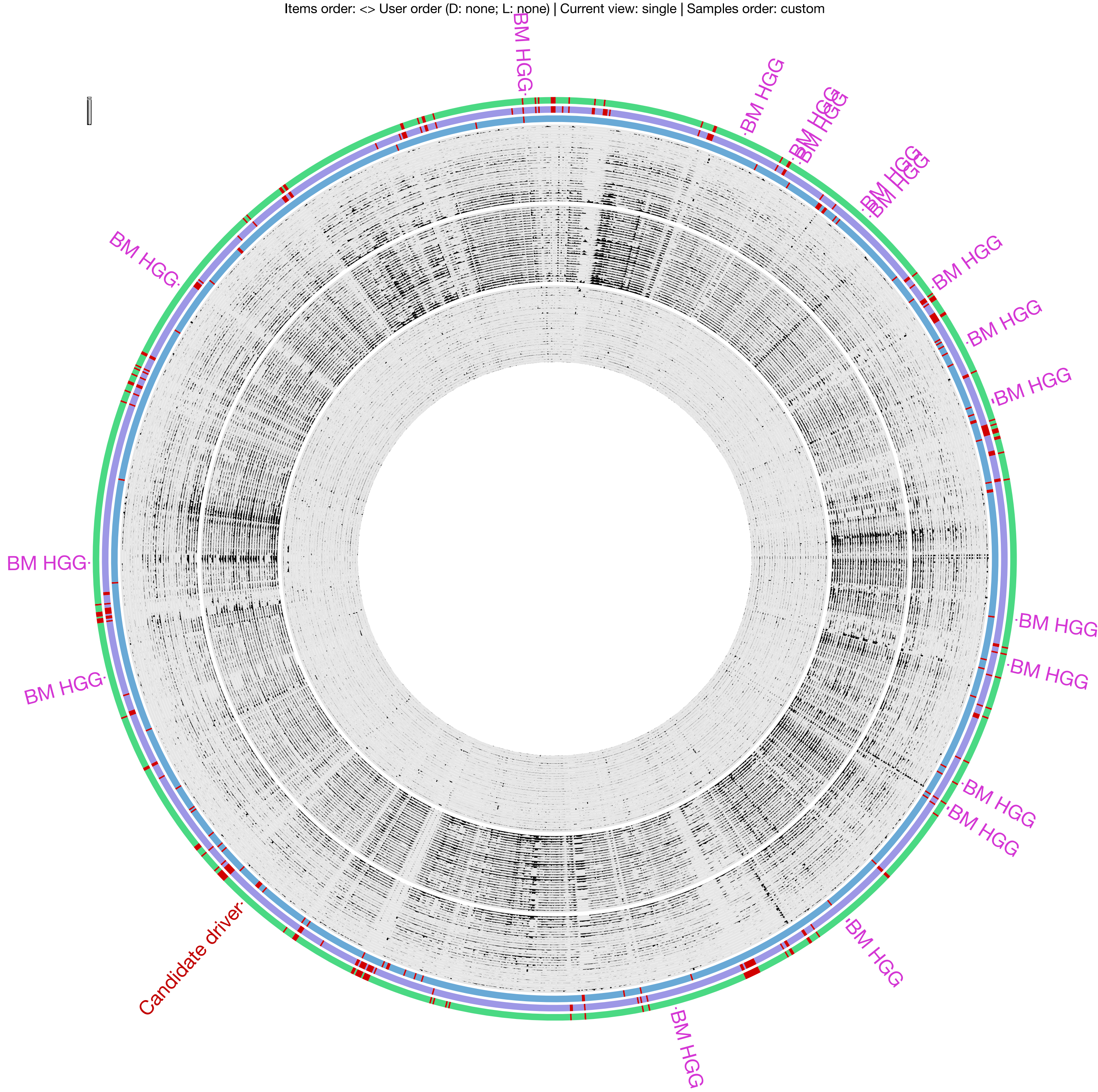


TD detection DETECTED (1721) NOT DETECTED (69)

BM detection DETECTED (1633) NOT DETECTED (157)

SUPP detection DETECTED (1667) NOT DETECTED (123)
