## Supplemental Methods for "Metapangenomics of the oral microbiome provides insights into habitat adaptation and cultivar diversity": Supplemental_Methods.html

Oral Metapangenomics


### Oral Metapangenomics

###### Daniel Utter

###### 13 January 2020

#### Overview

This is the long-form, narrative version of the metapangenomic methods used for our study Metapangenomics of the oral microbiome provides insights into habitat adaptation and cultivar diversity. Our goal in writing this is to make the workflow be as transparent and reproducible as possible, to fully describe the parameter and other methodological choices with data, and also to provide a step-by-step workflow for interested readers to adapt our methods to their own needs in systems within or outside of the oral microbiome. Most of the code here is based on Tom Delmont and A. Murat Eren’s *Prochlorococcus* metapangenome workflow, and we are deeply indebted to their project and their dedication to sharing reproducible workflows.

Our project investigated the environmental representation of the various bacterial pangenomes to learn more about the ecology and evolution of microbial biogeography. We were particularly interested in identifying and understanding the population structure of bacteria within the oral cavity, and generating as short as possible lists of candidate genes or functions that may be involved in the observed population structure, using the healthy human mouth as a model system. The mouth is well-suited to such studies of habit adaptation, as it consists of many distinct sites spatially connected by saliva but each with characteristically distinct community assemblages. We focused on three oral sites (hereafter ‘habitats’ to avoid ambiguity) - tongue dorsum (TD), buccal mucosa (BM), and supragingival plaque (SUPP). Many resident oral bacterial taxa have many high-quality genomes from cultured isolates, so we collected previously published genomes and combined this information with metagenomes from the Human Microbiome Project to address our goals.

This workflow shows the specific code used for *Haemophilus parainfluenzae*, and changing a few variable names will reproduce the *Rothia* analyses as well.

##### Organization & Workflow

Each oral habitat (tongue dorsum, TD; buccal mucosa, BM; supragingival plaque, SUPP) was processed independently in parallel. We used batch scripts and job arrays for this, and we include header information to provide context for the computational requirements for each step. Analyses for each oral habitat referred to the same annotated contigs database (holding the same genome nucleotide sequences), to which a different set of metagenomes were mapped, and downstream files were kept distinct by appending site identifiers as suffixes, e.g. `-bm`, to file names. This way, everything occurs in the same directory but with minimal duplication of scripts. Ultimately, three independent pangenomes were created, one from each habitat, and then exported to be overlaid onto a single pangenome figure. Following is a visual example of how select files and directories were shown:

```
+-- hpara_metapan
|   +-- 01_genContigsDB.sh
|   +-- 02_btmapHMP.sh
|
|   +-- Haemophilus-isolates.fa
|   +-- Haemophilus-isolates-CONTIGS.db
|   +-- Haemophilus-isolates-td-PAN
|   |   +-- Haemophilus-isolates-td-PAN.db
|
|   +-- hmp_all_td
|   |   +-- SRS_*fastq.gz
|   +-- hmp_all_bm
|   |   +-- SRS_*fastq.gz
|   +-- hmp_all_supp
|   |   +-- SRS_*fastq.gz
|
|   +-- bt_mapped_Haemophilus-isolates-td
|   |   +-- SRS_*.bam
|   |   +-- SRS_*.bai
```

**Side note:** We ran all of the anvi’o workflows on Harvard University’s Canon cluster, which uses the Slurm scheduler. We have kept the Slurm header information specifying the cluster parameters requested, since this includes the parallelization information. Also, this gives some context for memory and time requirements for replicability.

#### Step 1 - Collecting and annotating genomes

##### Getting the genomes from NCBI

Genomes were programmatically downloaded from NCBI RefSeq using the files found at ftp://ftp.ncbi.nlm.nih.gov/genomes/ASSEMBLY\_REPORTS/ as of 24 July, 2018. Note that not all genomes on NCBI are free of contamination, and so it is critical to inspect the quality of each genome before any final interpretations. Unfortunately, it is difficult to know which genomes have contamination issues prior to downloading, thus we downloaded and processed all, and then used the analytics produced in this workflow (coverage evenness across the genome, variability, genome size, GC content) to identify contaminants. For *H. parainfluenzae*, no genomes evidenced severe contamination, but for *Rothia*, a handful of genomes ultimately showed questionable gene content at later steps (discussed later) and so were removed, and the analysis was re-run with the reduced set of genomes.

Here is the ad hoc downloader script we wrote to programmatically download these. The NCBI assembly sheet, taxon to search for, and directory into which the genomes are downloaded are specified in the top 3 lines. Please note that this script renames the contig deflines to be A) simple and anvi’o compatible and B) contain the assembly sheet’s genus, species, and strain identifier, so that later the originally assigned taxonomy can be recovered. Since these genomes were originally downloaded, anvi’o now supports a more elegant workflow for downloading NCBI genomes; we include this code not as a recommendation of our ad hoc method over anvi’o’s method, but simply to detail how it was done:

```
#!/bin/bash

assemblyFile="assembly_summary_refseq.txt"
taxa="Haemophilus_parainfluenzae"
taxDir="Haemophilus_parainfluenzae"

mkdir ncbi_genomes
mkdir ncbi_genomes/$taxDir

for taxon in $taxa; do

# make sure whitespaces or lack thereof are ok
taxonSafe=$(echo $taxon | tr ' ' '_')
taxonGrep=$(echo $taxon | tr '_' ' ')

# assembly directory on the ncbi ftp site is the 20th column in assembly file
ftpDir=$(grep "$taxonGrep" $assemblyFile | awk -F"\t" '{print $20}')

cd ncbi_genomes/$taxDir

for dirPath in $ftpDir; do

# get just the ID
genomeID=$(echo "$dirPath" | sed 's/^.*\///g')
# get the path to the genomic fasta file
ftpPath="$dirPath/${genomeID}_genomic.fna.gz"
# make a friendly prefix - 'Gspe' from 'Genus species'
friendlyName=$(grep "$genomeID" ../../$assemblyFile | awk -F"\t" '{print $8}' | awk '{print $1 FS $2}' | tr -d '_,.-' | sed -E 's/^(.)[A-z0-9]* ([A-z0-9]{2}).*$/\1\2/')


# figure out the taxon + strain id from sheet to make deflines; 9 = strain,10=additional info
# this makes it simple to go from anvio contigs db to genome's original strain ID on NCBI
deflineName=$(grep "$genomeID" ../../$assemblyFile | awk -F"\t" '{print $8 " " $9}' | sed 's/strain=//' | tr ' -' '_' | tr -d '/=(),.')

genomeID="${friendlyName}_$genomeID"

# download & uncompress
wget $ftpPath -O $genomeID.fa.gz
gunzip $genomeID.fa.gz

# rename the deflines
awk -v defline="$deflineName" '/^>/{print ">" defline "_ctg" (++i)}!/^>/' $genomeID.fa >> temp.txt
mv temp.txt $genomeID.fa

done

done
```

##### Anvi’o contigs database

Now we combined the downloaded raw FASTAs into a single FASTA file to start the anvi’o workflow as follows:

```
cat ncbi_genomes/Haemophilus_parainfluenzae/*fa > Haemophilus-isolates.fa
```

###### Master script to prepare the contigs

At this point, an anvi’o contigs database is then generated from the single combined FASTA file, for which we invoked anvi’o’s HMM profiler, launched another script to annotation gene sequences, and made a bowtie2 reference database out of the original FASTA. As part of the `anvi-gen-contigs-database` command, Prodigal was run to call open reading frames (ORFs). We did this with the following script (we called it `01_genContigsDB.sh`). Note that this script submits two other scripts - specifically, `99_geneFunctions.sh` which runs the annotation script, and `02_btmapHMP.sh` which starts the next step.

```
#!/bin/bash
#SBATCH -N 1 #1 node
#SBATCH -n 6 # 1 cores from each
#SBATCH --contiguous
#SBATCH --mem=12G #per node
#SBATCH -t 0-12:00:00
#SBATCH -p shared
#SBATCH --job-name="HaemCont"
#SBATCH -o odyssey_cont.out
#SBATCH -e odyssey_cont.err
#SBATCH --mail-type=END

genus="Haemophilus"

# need to activate venv directly because node receiving job doesn't like bashrc aliases
source ~/virtual-envs/anvio-dev-venv/bin/activate

# clean up fasta deflines in contig file at the start for smooth downstream
anvi-script-reformat-fasta -o $genus-isolates-CLEAN.fa --simplify-names -r $genus-isolates-contigIDs.txt $genus-isolates.fa

mv $genus-isolates-CLEAN.fa $genus-isolates.fa

# make the contig db
anvi-gen-contigs-database -f $genus-isolates.fa -n $genus -o $genus-isolates-CONTIGS.db

# get bacterial single-copy gene, 16S, info from hmm collections
anvi-run-hmms -c $genus-isolates-CONTIGS.db --num-threads 6

# get the functional annotation started
sbatch 99_geneFunctions.sh

# make bt of contigs for mapping later
module load samtools/1.5-fasrc01 bowtie2/2.3.2-fasrc01
bowtie2-build $genus-isolates.fa $genus-isolates

# start next step
sbatch 02_btmapHMP.sh
```

The purpose of the HMM models run here with `anvi-run-hmms` is to identify bacterial single-copy core genes in each genome. From this information, the relative completion and redundancy of each genome can be estimated from the number of expected single copy genes found once or twice, respectively. This step is critical, because in addition to other ways of estimating assembly quality (number of contigs, length of assembly, and number of genes) it allows us to investigate whether genomic similarities result from similarly poor genome assemblies, or whether genome assembly is comparable and thus any similarities (or differences) are due to potentially biological reaasons.

###### Functional annotation

We used Interproscan to get functional annotation from a variety of databases to ensure that we were not overly sensitive to the lacking of any one functional database. The core command consists of:

```
sh PATH/TO/INTERPRO/interproscan-5.36-75.0/interproscan.sh -i genes.faa -o genes.tsv -f tsv --appl TIGRFAM,Pfam,SUPERFAMILY,ProDom
```

However, for cases like *Rothia* with 67 genomes’ worth of genes, a non-parallel workflow took too long, so we made a simple parallelization. Here is the annotation script we used as `99_geneFunctions.sh`:

```
#!/bin/bash

prefix="Haemophilus-isolates"

# activate anvio if not already
source ~/virtual-envs/anvio-dev-venv/bin/activate

# export amino acid sequences
anvi-get-sequences-for-gene-calls --get-aa-sequences --wrap 0 -c $prefix-CONTIGS.db -o $prefix-gene-calls-aa.faa

batchDir="$prefix-gene-calls-batches"
mkdir $batchDir

# chop file up into 25k sequences per file
split -l 50000 -d -a 4 $prefix-gene-calls-aa.faa $batchDir/$prefix-gene-calls-aa.faa-

# figure out how many batches we made
hiBatch=$(ls $batchDir/*.faa-* | tail -1 | sed 's/^.*-0*\([1-9]*\)\(.$\)/\1\2/')
numBatches=$(($hiBatch + 1))

# set up the file to concatenate into
echo -e "gene_callers_id\tsource\taccession\tfunction\te_value" > interpro-results-fmt-$prefix.tsv

echo "#!/bin/bash
#SBATCH -N 1 #1 nodes of ram
#SBATCH -n 12 # 12 cores from each
#SBATCH --contiguous
#SBATCH --mem=12G #per node
#SBATCH -t 0-6:00:00
#SBATCH -p shared
#SBATCH --array=0-$hiBatch%$numBatches
#SBATCH --job-name='interpro-%a'
#SBATCH -o odyssey_anviIP_%a.out
#SBATCH -e odyssey_anviIP_%a.err
#SBATCH --mail-type=END

# format job array id to match the split output
taskNum=$(printf %04d $SLURM_ARRAY_TASK_ID)

batch=$batchDir/$prefix-gene-calls.faa-$taskNum

module load jdk/1.8.0_172-fasrc01

./../../interproscan-sh/interproscan-5.36-75.0/interproscan.sh -i $batch -o ${batch}.tsv -f tsv --appl TIGRFAM,Pfam,SUPERFAMILY,ProDom,Gene3D

# format it manually to be safe 
cat ${batch}.tsv | awk -F\"\t\" '{print $1 FS $4 FS $5 FS $6 FS $9}' >> interpro-results-fmt-$prefix.tsv

" > ip-$batch.sh && sbatch ip-$batch.sh && rm ip-$batch.sh
```

We invoked `01_genContigsDB.sh`, which read the contig FASTAs into an anvi’o contigs database and then called the functional annotation script, which wrote the file `interpro-results-fmt-Haemophilus-isolates.tsv` containing the annotation information. After everything here finished, we incorporated the functional annotation into the contigs database with the command

```
anvi-import-functions -c Haemophilus-isolates-CONTIGS.db -i interpro-results-fmt-Haemophilus-isolates.tsv
```

Separately, we added NCBI COG annotations, to have larger functional categories to enable broader comparisons, with the following script:

```
#!/bin/bash
#SBATCH -N 1 #1 nodes of ram
#SBATCH -n 20 # 1 cores from each
#SBATCH --contiguous
#SBATCH --mem=12G #per node
#SBATCH -t 1-12:00:00
#SBATCH -p shared
#SBATCH --job-name="COGhp"
#SBATCH -o odyssey_COG-hp.out
#SBATCH -e odyssey_COG-hp.err
#SBATCH --mail-type=END

# activate anvio
source ~/virtual-envs/anvio-dev-venv/bin/activate

module load ncbi-blast/2.10.0+-fasrc01

prefix="Haemophilus-isolates"

anvi-run-ncbi-cogs -c $prefix-CONTIGS.db --search-with blastp -T 18
```

The resultant contigs database for both *H. parainfluenzae* and *Rothia* can be found in this FigShare dataset.

#### Step 2 - Mapping

##### Data acquisition - HMP Metagenomes

Raw short-read metagenomic data from the Human Microbiome Project (HMP) (HMP, 2012; Lloyd-Price et al., 2017) was downloaded for the tongue dorsum (TD, n = 188), buccal mucosa (BM, n = 169), and supragingival plaque (SUPP, n = 194) sites using HMP data portal at https://portal.hmpdacc.org/. Paths for each sample were downloaded via the HMP portal and uploaded to Cannon, from which we ran an ad-hoc script to download all of them. We used the S3 method as it was the most reliable. Here is the script we used for downloading the buccal mucosa metagenomes:

```
#!/bin/bash
#SBATCH -N 1 #1 nodes of ram
#SBATCH -n 1 # 1 cores from each
#SBATCH --contiguous
#SBATCH --mem=2G #per node
#SBATCH -t 0-6:00:00
#SBATCH -p shared
#SBATCH --array=0-33%34
#SBATCH --job-name="dlHMP"
#SBATCH -o odyssey_downloadHuman-%a.out
#SBATCH -e odyssey_downloadHuman-%a.err
#SBATCH --mail-type=NONE

# cat buccal_manifest.txt | split -l 6 -d -a 3 - hmpBatch_

# format to match what we made
taskNum=$(printf %03d $SLURM_ARRAY_TASK_ID)

batch="hmpBatch_$taskNum"

# activate anvio
source ~/virtual-envs/anvio-dev-venv/bin/activate

#loop through each sample in this batch taking only the aws location
for hmp in $(sed 's/^.*s3/s3/; s/bz2.*$/bz2/' $batch); do

sh -c 'aws s3 cp "$0" .' $hmp
tar -xjf $(echo "$hmp" | sed 's,^.*/,,g')

done
```

Here, the `buccal_manifest.txt` file was the saved text file of the HMP DACC cart for BM metagenomes, after removing the header line. Prior to running this script, we ran the commented-out line `cat buccal_manifest.txt | split -l 5 -d -a 3 - hmpBatch_` as a one-liner at the command prompt to split the manifest into 5-metagenome batches that the array would then process, and we manually changed the array parameters `--array=0-33%34` to match the number of batches this made.

This same process was performed for tongue dorsum (TD), buccal mucosa (BM), and supragingival plaque (SUPP), each of which in its own directory, e.g., `hmp_all_bm/`.

After downloading and uncompressing, the R1 and R2 reads were in a subdirectory for each metagenome, so we moved the R1/R2 pairs out of these subdirectories leaving the junk singleton reads behind (e.g. `mv */*.1.fastq .`), then deleted the downloaded \*bz2 files and the directories.

##### Recruiting HMP metagenomes to the contigs

After downloading the metagenomes as described above, we competitively recruited the HMP metagenomes onto the exact contigs included in the contigs database using bowtie2 (Langmead & Salzberg, 2012) with default parameters (`--sensitive`). `--sensitive` mode was chosen as we wanted a balance between strict matching, but not too strict as we want to identify different but closely-related *H. parainfluenzae* and *Rothia* populations living in the mouth. Since all the reference genomes originate from cultivated isolates, we cannot assume that they perfectly match natural, uncultivated populations in the mouth, and so we must allow for some differences in mapping.

Metagenomes from each oral habitat (TD, BM, SUPP) were mapped onto separate but identical contigs databases. We did this with one script that made a separate job array for each habitat:

```
#!/bin/bash

# 20171016 - assign gene fams by interproscan
prefix="Haemophilus-isolates"

# this sets up the site suffixe to iterate through in parallel
sites=(bm td supp)

# iterate through each habitat - this will create a custom script for each habitat that is an array over all that sites metagenomes
for site in ${sites[*]}; do

mkdir bt_mapped_${prefix}_$site
mkdir batches_$site

# make batches for each sites samples
ls --color=none hmp_all_$site/*1.fastq | split -l 8 -d -a 3 - batches_$site/$site-

# figure out the upper bound of the array for slurm
hiBatch=$(ls batches_$site/$site-* | tail -1 | sed 's/^.*-0*\([1-9]*\)\(.$\)/\1\2/')
numBatches=$(($hiBatch + 1))

echo "#!/bin/bash
#SBATCH -N 1 #1 nodes of ram
#SBATCH -n 12 # 1 cores from each
#SBATCH --contiguous
#SBATCH --mem=18G #per node
#SBATCH -t 0-06:00:00
#SBATCH -p shared
#SBATCH --array=0-$hiBatch%$numBatches
#SBATCH --job-name='bt-$site'
#SBATCH -o odyssey_bt-$site-%a.out
#SBATCH -e odyssey_bt-$site-%a.err
#SBATCH --mail-type=NONE

# format this array element id to match batch name
taskNum=\$(printf %03d \$SLURM_ARRAY_TASK_ID)

module load samtools/1.5-fasrc02 bowtie2/2.3.2-fasrc02 xz/5.2.2-fasrc01

# set what batch we are on
batch=\"batches_$site/$site-\$taskNum\"

# loop through metagenomes to map in this batch
for FQ in \$(cat \$batch); do 

# FQ was the R1 path so extract the R2 path from it
r2=\$(echo \"\$FQ\" | sed 's/.1.fastq/.2.fastq/')

bowtie2 -x $prefix -1 \$FQ -2 \$r2 --no-unal --threads 12 | samtools view -b - | samtools sort -@ 12 - > \$(echo \"\$FQ\" | sed 's/hmp_all/bt_mapped_$prefix/; s/\.1.fastq.*$/.bam/')

# index as next anvio step requires indexed bams
samtools index \$(echo \"\$FQ\" | sed 's/hmp_all/bt_mapped_$prefix/; s/\.1.fastq.*$/.bam/')

done
" > bt-$site-array.sh && sbatch bt-$site-array.sh && rm bt-$site-array.sh # save this file and submit it then delete for housekeeping

sleep 30

done
```

This script is `02_btmapHMP.sh` and was run directly from `01_genContigsDB.sh` (Step 1)

###### A short statement on competitive mapping and its interpretation

Each sample’s short reads were mapped against all genomes simultaneously for that taxon; thus, bowtie2 matched each read to the best-matching genomic locus, randomly choosing between multiple loci if they were equally best. Thus, coverage at highly conserved regions is affected by the total population abundance in that sample, while the coverage at variable loci reflects that particular sequence variant’s abundance. As such, this competitive recruitment approach can discriminate between highly dissimilar reference sequences, as well as apportion the diversity encompassed by a natural population onto the closest member(s) of the reference genome set.

Regions of identical nucleotide similarity between reference genomes, relative to the metagenome, then recruit an approximately equivalent number of reads, reflecting their proportion of the total population abundance in that sample. However, regions with polymorphic sites allow for discrimination between genomes, based the variability of sequences in the pangenome / database.

At the extreme, highly conserved genes or regions (such as some regions of the 16S rRNA gene) are so conserved that identical and nearly identical sequences can be found in closely related taxa. Similarly, mobile genetic elements like phage or transposons can also occur outside the target population. Such sites receive much more coverage than their surrounding genome. For this reason, the coverage of each gene in each genome was inspected 1) to ensure that coverage estimates are not critically biased by conserved or mobile genes and 2) identify potential contaminant genes in the reference genomes, whether human or otherwise. This section continues to describe the methods for obtaining the gene-level mapping data; the section “Visualizing mapping results at the per-gene, per-sample level” describes the interpretation of gene-level mapping patterns from these data.

#### Step 3 - Profiling the metagenome recruitment

After mapping, we profiled each sample’s coverage into an anvi’o profile database and then merged them all. As for the mapping step, we wrote one master script with the same overall architecture as before.

Individual sample profiling code:

```
#!/bin/bash

# 20171016 - assign gene fams by interproscan
prefix="Haemophilus-isolates"

# need to activate venv directly because node receiving job order doesn't play with bashrc aliases

# habitats to iterate over
sites=(bm td supp)

# for each habitat we generate a custom job array
for site in ${sites[*]}; do

mkdir profs_mapped_${prefix}_$site

# make batches for each sites samples but leaving commented off as the bowtie2 batches work here too
#ls --color=none hmp_all_$site/*1.fastq | split -l 8 -d -a 3 - batches_$site/$site-

# find high number of batches we made before
hiBatch=$(ls batches_$site/$site-* | tail -1 | sed 's/^.*-0*\([1-9]*\)\(.$\)/\1\2/')
numBatches=$(($hiBatch + 1))

echo "#!/bin/bash
#SBATCH -N 1 #1 nodes of ram
#SBATCH -n 12 # 1 cores from each
#SBATCH --contiguous
#SBATCH --mem=18G #per node
#SBATCH -t 0-08:00:00
#SBATCH -p shared
#SBATCH --array=0-$hiBatch%$numBatches
#SBATCH --job-name='pro-$site'
#SBATCH -o odyssey_prof-$site-%a.out
#SBATCH -e odyssey_prof-$site-%a.err
#SBATCH --mail-type=NONE

taskNum=\$(printf %03d \$SLURM_ARRAY_TASK_ID)

source ~/virtual-envs/anvio-dev-venv/bin/activate

batch=\"batches_$site/$site-\$taskNum\"

# for each fastq that was mapped
for FQ in \$(cat \$batch); do

# recreate the name of the bam from the name of the fastq
bam=\$(echo \"\$FQ\" | sed 's/hmp_all/bt_mapped_$prefix/; s/\.1.fastq.*$/.bam/')

# generate a profile
anvi-profile -i \$bam -c $prefix-CONTIGS.db -W -M 0 -T 12 --write-buffer-size 500 -o \$(echo \"\$bam\" | sed 's/bt/profs/; s/.bam//')

done
" > prof-$site.sh && sbatch prof-$site.sh && rm prof-$site.sh

sleep 25
done
```

Note we set the -M flag, which controls the minimum contig length to analyze, to zero. From the anvi’o documentation, `-M` parameter has a different default:

> Minimum length of contigs in a BAM file to analyze … we chose the default to be 2500 [nt]

Since a few genomes we analyzed contained contigs with fewer than 2500 nt, we elected to maintain all contigs, rather than dropping contigs or genomes. The anvi’o default of 2.5 kb may be for the purpose of meaningful tetranucleotide frequency comparisons during metagenomic bin refinement. We justify our choice as we are not refining bins but rather working with existing genomes from nominally axenic isolates. However, as short contigs are more likely to contain contaminant genes than longer genes, this choice opens up the possibility of including contaminant contigs. Since our approach ultimately allows inspection of the per-gene coverage of each genome, we can identify contaminant genes later since these genes almost always have aberrant coverage profiles relative to the surrounding genome or might be expected from their identity (e.g., >1000x coverage of genes that are neither ribosomal RNAs, transposons, nor phage when rest of the genome has 0-10x coverage). Details of this inspection are discussed in the section “Vizualizing mapping results at the per-gene, per-sample level.”

Another key factor is that we used the same contigs database for all the profiles. In other words, we made one `*-CONTIGS.db` at the start, and then mapped hundreds of metagenomes onto the same FASTA that generated it, and now have made over a hundred each of TD, BM, and SUPP metagenome profiles for this one set of contigs.

#### Step 4 - Pangenome and metapangenome construction

This step incorporates multiple related sub-steps. The overall workflow is to merge each habitat’s profiles into a single merged profile database, then store the information about which contigs and genes belong to which genomes since that hasn’t been done yet. Then, with this information we can make a pangenome. Once we have this, since we have the 3 profile databases that contain the information about coverage, SNPs, etc. for each metagenome sample, and this information corresponds to the genes in the shared CONTIG.db, we can overlay that metagenomic data onto the pangenome to have a metapangenome. This was all accomplished in one script, as follows:

```
#!/bin/bash

# 20171016 - assign gene fams by interproscan
prefix="Haemophilus-isolates"

# make a two-column table of which contigs go with which genomes for later, easy since original contigs had this already
sed 's/_ctg[0-9]*$//' $prefix-contigIDs.txt > $prefix-renaming-manual.tsv

# list of habitats to iterate over
sites=(td bm supp)
mcl=10
#mcls=(8 10 12) # for varying mcl
#site="td" # for varying mcl

for site in ${sites[*]}; do
#for mcl in ${mcls[*]}; do  # for varying mcl

echo "#!/bin/bash
#SBATCH -N 1 #1 node
#SBATCH -n 1 
#SBATCH --contiguous
#SBATCH --mem=60G #per node
#SBATCH -t 1-18:00:00
#SBATCH -p shared
#SBATCH --job-name=\"fin-$site-$indiv\"
#SBATCH -o odyssey_metapan-$site-$indiv.out
#SBATCH -e odyssey_metapan-$site-$indiv.err
#SBATCH --mail-type=END

source ~/virtual-envs/anvio-dev-venv/bin/activate
module load mcl/14.137-fasrc01 ncbi-blast/2.6.0+-fasrc01 #blast-2.6.0+-fasrc01

anvi-merge profs_mapped_${indiv}_$site/*/PROFILE.db -c $indiv-CONTIGS.db -o $indiv-$site-MERGED -W

anvi-import-collection -c $indiv-CONTIGS.db -p $indiv-$site-MERGED/PROFILE.db -C Genomes --contigs-mode $indiv-COLLECTION-MAPPER.txt
anvi-summarize -c $indiv-CONTIGS.db -p $indiv-$site-MERGED/PROFILE.db -C Genomes -o $indiv-$site-SUMMARY

echo 'done summarizing'

./anvi-script-gen-internal-genomes-table.sh $indiv-$site

sed \"s/$indiv-$site-CONTIGS.db/$indiv-CONTIGS.db/g\" $indiv-$site-internal-genomes-table.txt > tmp
mv tmp $indiv-$site-internal-genomes-table.txt

anvi-gen-genomes-storage -i $indiv-$site-internal-genomes-table.txt -o $indiv-$site-GENOMES.db
anvi-pan-genome --mcl-inflation $mcl -o $indiv-$site-$mcl-PAN -g $indiv-$site-GENOMES.db -n $indiv-$site-$mcl --use-ncbi-blast -T 12

anvi-meta-pan-genome -i $indiv-$site-internal-genomes-table.txt -g $indiv-$site-GENOMES.db -p $indiv-$site-$mcl-PAN/*PAN.db
" > finish-$site.sh && sbatch finish-$site.sh && rm finish-$site.sh

sleep 10

done
```

The `anvi-pan-genome` family of commands requires a specially-formatted table to tell anvi’o where to look. This is the helper script `anvi-script-gen-internal-genomes-table.sh` referred to in the above script that we wrote to collect all this info:

```
#!/bin/bash

taxonPrefix=$1
collection="Genomes"

echo -e "name\tbin_id\tcollection_id\tprofile_db_path\tcontigs_db_path" > $taxonPrefix-internal-genomes-table.txt

for bin in $(ls $taxonPrefix-SUMMARY/bin_by_bin); do

echo -e "$bin\t$bin\t$collection\t$taxonPrefix-MERGED/PROFILE.db\t$taxonPrefix-CONTIGS.db" >> $taxonPrefix-internal-genomes-table.txt

done
```

This wrote the needed info as `*-internal-genomes-table.txt` that was used for the pangenome steps.

At this point, for our *H. parainfluenzae* specific case shown here, we have one directory with a single contigs db called `Haemophilus-isolates-CONTIGS.db` and three merged profile databases like `Haemophilus-isolates-bm-MERGED/PROFILE.db` with coverage, detection (fraction of gene or genome with >=1x coverage) and metagenome nucleotide variant data from all the metagenomes for that habitat, and three genome databases like `Haemophilus-isolates-bm-GENOMES.db` and three pangenome databases like `Haemophilus-isolates-bm-10-CONTIGS.db` where the number in the file name reports the MCL inflation factor used (explained more in the next section.)

The `*GENOMES.db` and `*PAN` databases can be found in this FigShare dataset.

###### Pangenome creation methods details

The pangenome was calculated with the anvi-pan-genome command. We wanted to compare homologous genes between all genomes (here, *H. parainfluenzae*), so we first needed to cluster the observed genes (technically, ORFs) into homologous units – what anvi’o and we refer to as **gene clusters** since they are the result of clustering gene sequences in amino acid space. For more background, please see the step-by-step anvi’o documentation and explanation here.

Briefly, this clustering takes advantage of the fact that pairwise amino acid similarity is not continuous as phylogenetic relatedness decays, but rather decreases in a stair step pattern. That is, differences between any pair of true homologs within a given species or genus should be much smaller than the difference between one of those genes and any non-homologous gene. However, since different gene families may evolve and diversify at different rates, the exact similarity threshold that delineates homologs may vary.

Visually, gene pairwise similarities might look like the following for four theoretical groups of genes:

And from this plot, it would be easy to identify A) that there are 4 groups of genes (colored dots), and B) which genes belong in which group. In this example, there is a fairly uniform step-size, and groups could be defined either based on a drop in similarity or by some set threshold (e.g. > 92% similarity).

On the other hand, another plausible scenario could also look like this:

This plotted scenario highlights how an absolute threshold of similarity is uninformative for different genes that may evolve more slowly or quickly relative to each other. But, the drop in similarity between genes distinguishes the groups, although one or two edge-case genes may be misplaced.

Thus, the most general-purpose approach is to find the drop in similarity between all gene pairs and use this information to define groups of genes. So, if one imagines each gene as a node on a network, and each node is connected to other nodes by edges that are amino acid sequence similarity, homologous units would appear as groups of nodes (genes) connected more tightly than random, such that if a random walk through the network traversed one node in that group, it would mostly likely traverse other nodes in that cluster.

And this is how the algorithm MCL (van Dongen & Abreu-Goodger, 2012) works when applied to pairwise gene similarities.

We used BLASTP (Altschul et al., 1990) to compute amino acid-level similarities between all possible ORF pairs after alignment with MUSCLE (Edgar, 2004). Too-weak matches were culled by employing the `--minbit` criterion with the default value of 0.5, to minimize feeding MCL irrelevant similarities - by calculating all possible pairwise similarities, we introduce nonsensical comparisons, e.g., DnaK to TonB, that can be disregarded by their poor alignment quality prior to MCL. Briefly, `--min-bit 0.5` requires the BLAST bitscore between any two genes to be at least half of the maximum bitscore allowed by the shortest of the two sequences. MCL then uses these pairwise identities to group ORFs into gene clusters, putatively homologous gene groups. MCL uses a hyperparameter (inflation, `--mcl-inflation`) to adjust the clustering sensitivity, i.e., the tendency to split clusters. For the nominally single-species *H. parainfluenzae* pangenome we used `--mcl-inflation 10`; for all other pangenomes which were genus-level, we used `--mcl-inflation 6`.

To test the sensitivity of our pangenome results, specifically the number and emerging core/accessory patterns of gene clusters, we varied the MCL inflation value by ±2 for each pangenome, and compared the number of gene clusters produced as well as comparing the core/accessory patterns. Computationally, this was done with the script above but by uncommenting the `mcls=(8 10 12)` line, fixing the `site` variable to a specific site (such as `site='bm'`), and setting the for loop to iterate through the `mcls`. The total number of gene clusters changed by only -2.3% to 1.6% for *Rothia* and -0.4% to 0.5% for *H. parainfluenzae* (exact values in Supplementary Table 1), and the overall patterns and proportional size of the core/accessory genome appeared nearly identical ( Additional File 11 ).

###### Metapangenome detailed assumptions and specifics

One of the main goals of overlaying metagenome data onto a pangenome is to understand which genes in the pangenome are common in environmental populations and which are rare. To make this assessment, two criteria are needed: First, a criterion for the genome to establish whether the genome occurs at all in the metagenome(s) (i.e., whether the metagenomes are appropriate to study the genome and its genes), and second, a criterion for each gene to establish whether the gene is ‘core’ or ‘accessory’ to the population sampled by the metagenome.

We set the `--min-detection` parameter in `anvi-meta-pan-genome` to the default value of `0.5`. Thus, if half or more of the total nucleotides in a genome are not covered by at least one metagenome for that habitat, the program skips trying to calculate any results and returns a value of `NA` instead. This threshold protects us from cases where we could be looking at coverage disparities between a gene and its genome, but it is simply a result of the fact that the genome is not represented in this habitat. `0.5` is ultimately an arbitrary but useful threshold, as the detection of a handful of genes/regions of a genome can certainly occurs from spurious causes, but it is difficult to imagine a plausible scenario where greater than half of a genome receives coverage from exclusively spurious sources. On the other hand, intermediate-but-too-low detection, such as between 0.1 and 0.4, most plausibly originate from scenarios where a population similar to the reference genome exists, but at such a low abundance in the sample that the sequencing depth was too low to adequately sample that population. Thus, only analyzing genomes with at least half of the genome receiving 1x coverage ensures that we consider only genome/metagenome combinations where the sequencing depth is sufficient and where the metagenome likely contains at least half of a genome’s genes.

Here, the `--fraction-of-median-coverage 0.25` parameter means that we set the threshold between ‘**environmental core genes**’ or ‘**environmental accessory genes**’ (ECG and EAG) for each gene’s median coverage across a habitat’s metagenomes to be 0.25 of the genome’s corresponding coverage across those same samples. For instance, *H. parainfluenzae* T3T1 obtained a median coverage of 12X across the 188 TD metagenomes, implying that since the median TD metagenome covered T3T1 12X, then a given gene in the T3T1 genome obtaining a median coverage of 1.2X from those same 188 metagenomes would be considered environmental accessory in TD. However, if a different T3T1 gene had a median coverage of 4.8X or 27.6X, then that gene would be environmental core in TD.

Like the minimum detection threshold, the environmental core/accessory threshold is arbitrary but useful. Biologically, a genome occurring in approximately half of the members of a population is certainly not core to the population, and so halving that threshold (to 0.25) offers extra caution to not falsely define environmentally accessory genes. However, the specific threshold value chosen to define environmental core/accessory is largely unimportant, as the majority of genes are either completely detected in the majority of metagenomes, or are completely undetected in most metagenomes (see next section).

By defining the concept of environmentally core/accessory relative to genomic coverage, we can scale our expectation for finding a given gene in a habitat based on the genome. We do this so that when our metric reports that a gene is ‘environmental accessory’ in a habitat, we know that the metric is not simply labelling every gene from a low-abundance genome as environmental accessory, but rather only the genes that are much less covered in its habitat than expected based on coverages of the surrounding genome Thus, the coverage of the surrounding genome offers an expectation for how abundant the population is, and the identification of core or accessory is relative to the abundance of that population. So, this determination of environmental core/accessory is a proxy for whether a gene sequence is at a notably lower frequency in the environment than we presume its background genome to have (which could be a result of selection). Importantly, a gene being environmentally accessory has a much more confident interpretation than it being environmentally core. That is, a gene cannot be essential to a cell’s survival in a habitat and not occur in every cell living in that habitat, barring certain public-goods scenarios. On the other hand, there are many non-selective circumstances that could lead to a gene occurring as many times as there are cells without being essential.

Relating the gene’s median coverage to the genome’s median coverage also helps even out noise in gene-level coverages from non-specific mapping (falsely inflating coverage) or from having highly similar DNA sequences in the bowtie2 reference database competing for the same metagenome reads (lowering overall coverage).

###### Precise choice of ECG/EAG threshold has little impact

We wished to assess whether various `--fraction-of-median-coverage` thresholds would impact designation of genes as environmental core or environmental accessory. To this end, we investigated the distribution of different and more conservative measurement, detection (the fraction of a gene receiving any coverage at all), across metagenomes for each gene. We chose this approach, as inspection of gene-level coverages by metagenomes for each genome (discussed later) revealed that qualitatively, most genes appeared to be abundant and completely detected in all or most metagenomes and were environmentally core, while environmentally accessory genes were typically completely absent from the majority of metagenomes. Thus, if the distribution of gene detection is bimodal, any coverage-defined value for the ECG/EAG threshold, whether 0.2, 0.25, 0.4 of the parent genome’s median coverage, would be largely equivalent, since genes that are EAG are EAG because no part of the gene received coverage in the majority of metagenomes.

We collected the gene-level detection and coverage values from each metapangenome as follows:

```
#!/bin/bash
#SBATCH -N 1 #1 nodes of ram
#SBATCH -n 1 # 12 cores from each
#SBATCH --contiguous
#SBATCH --mem=1G #per node
#SBATCH -t 0-01:00:00
#SBATCH -p shared
#SBATCH --job-name="finish"
#SBATCH --mail-type=END

# 20171016 - assign gene fams by interproscan
indiv="Haemophilus-isolates"

sites=(td bm supp)

for site in ${sites[*]}; do

echo "#!/bin/bash
#SBATCH -N 1 #1 nodes of ram
#SBATCH -n 1 # 1 cores from each
#SBATCH --contiguous
#SBATCH --mem=80G #per node
#SBATCH -t 0-12:00:00
#SBATCH -p shared
#SBATCH --job-name=\"fin-$site-$indiv\"
#SBATCH -o odyssey_singlemetapan-$site-$indiv.out
#SBATCH -e odyssey_singlemetapan-$site-$indiv.err
#SBATCH --mail-type=END

source ~/virtual-envs/anvio-dev-venv/bin/activate

anvi-export-gene-coverage-and-detection -p $indiv-$site-MERGED/PROFILE.db -c $indiv-CONTIGS.db -O $indiv-$site

" > metapan-$site.sh && sbatch metapan-$site.sh  && rm metapan-$site.sh

sleep 10

done
```

We downloaded those output data, along with the genome-level detection, already calculated and located in the `-*SUMMARY/bins_across_samples/detection.txt` directory associated with each `*MERGED/PROFILE.db`. This detection table will allow us to only look at genes of genomes with a genome-wide detection of at least 0.5 in each habitat.

We also summarize the pangenome with `anvi-summarize` to have a convenient way to identify gene caller IDs associated with each genome.

```
site <- 'td'
prefix <- 'Haemophilus'

gc_summary <- read.csv(paste0(prefix,"-isolates-",td,"-10-SUMMARY/Haemophilus-isolates-",site,"-10_gene_clusters_summary.txt", sep="\t")
median_coverages <- read.csv(paste0(prefix,"-isolates-",td,"-10-SUMMARY/misc_data_layers/default.txt", sep="\t")

detection_genome <- read.csv(paste0(prefix,"-isolates-", site, "-SUMMARY/bins_across_samples/detection.txt"), sep = "\t")

detection_gene <- read.csv(paste0(prefix,"-isolates-", site, "-GENE-DETECTION.txt"), sep="\t")
coverage_gene <- read.csv(paste0(prefix,"-isolates-", site, "-GENE-COVERAGES.txt"), sep="\t")
```

Now that the data is prepared, we wrote a simple function to fetch the gene detection values for genes from a given genome for all metagenomes where that genome had a detection of at least 0.5, matching the `--min-detection` threshold.

```
extractGenes <- function(df_gene = coverage_gene, genome){
  genes_from_this_genome <- as.character(gc_summary$gene_callers_id[gc_summary$genome_name == genome])

  samples_genome_detected <- colnames(detection_genome[2:ncol(detection_genome)])[detection_genome[detection_genome$bins == genome, 2:ncol(detection_genome)] >= 0.5]
  filtered_gene_coverages <- df_gene[df_gene$key %in% genes_from_this_genome, samples_genome_detected]
  
  return(filtered_gene_coverages)
}
```

Then, we write a function to loop through all genomes, using the above helper function to extract gene detection from these values for each genome if that genome occurs in more than 1 metagenome. Then, it also extracts the gene coverage values and compares each gene to the genome’s median coverage to determine if that gene was ECG or EAG based on the 0.25 threshold.

```
library(reshape2)
library(ggplot2)
library(ggridges)

genomeList <- detection_genome$bins

getDetectionForECAGgenesForManyGenomes <- function(genomes=genomeList) {
  l <- list()
  for (genome in genomes) {
    all_ecag_detection <- extractGenes(df_gene = detection_gene, genome = genome)
    
    if (is.null(dim(all_ecag_detection))){
      print(paste0("Hi! Skipping ",genome," because it is detected in one or fewer samples"))
      next
    }
    
    all_ecag_coverage <- extractGenes(df_gene = coverage_gene, genome = genome)
    all_ecag_detection$gene <- rownames(all_ecag_detection)
    all_ecag_detection$ECAG <- "ECG"
    all_ecag_detection$ECAG[as.numeric(apply(all_ecag_coverage, 1, median)) / 
                              median_coverages[median_coverages$layers == genome, paste0(toupper(site),"_HMP_MEDIAN")] < 0.25] <- "EAG"
    
    all_ecag_long <- melt(all_ecag_detection)
    
    genome_pretty <- paste(gsub("^([A-Z])[a-z]*$","\\1", unique(unlist(strsplit(as.character(genome), "_")))), collapse = "_")
    all_ecag_long$genome <- genome_pretty
    
    l[[genome_pretty]] <- all_ecag_long
  }
  
  print(paste0("Got passed ", length(genomes), " genomes and prepped E[CA]G gene detection and ", length(l), " genomes were detected in >1 sample"))
  
  return(l)
}

many_ecag_genomes <- getDetectionForECAGgenesForManyGenomes()
```

```
## [1] "Got passed 33 genomes and prepped E[CA]G gene detection and 33 genomes were detected in >1 sample"
```

```
many_ecag_genomes_long <- melt(many_ecag_genomes)

many_ecag_genomes_long <- many_ecag_genomes_long[!is.na(many_ecag_genomes_long$value),]
many_ecag_genomes_long$genome[grep("dentocariosa_C6[BD]", many_ecag_genomes_long$genome)] <- gsub("dentocariosa","aeria", grep("dentocariosa_C6[BD]", many_ecag_genomes_long$genome, value = T))
```

So now we have the detection of each gene in each metagenome, along with whether that gene was determined to be ECG or EAG by having at least or less than (respectively) one-fourth of its parent genome’s median coverage.

```
ggplot(many_ecag_genomes_long, aes(x = value, group=gene, fill=ECAG)) + 
  scale_x_continuous(expand = c(0,0)) + scale_y_continuous(expand = c(0,0)) +
  facet_wrap(~genome) +
  labs(x = "Detection (fraction of gene with > 0X coverage)", y = "Number of metagenomes", fill="EAG or ECG") +
  geom_histogram(alpha=0.08, position = 'identity', bins=50) + theme_classic()
```

In this plot, each subplot shows the detection of genes from each genome. The x-axis is the fractional detection of a gene, and the y axis shows the number of metagenomes producing that detection for each gene. Note that the bars are translucent and overlaid, since there is one bar per gene for each 2% bin of detection values (since there are 50 bins for detection values from 0-1). From this plot, it is clear that there is a sharp increase in metagenomes producing the detection extremes relative to intermediate detections, i.e., genes with intermediate detection receive such intermediate detection in far fewer metagenomes than do genes completely detected or undetected. And, since genes with a detection of 0 by definition have no coverage, an exact fractional coverage threshold would have little importance for the lower extreme. However, estimating the relative likelihood of whether or not most genes fall into these extreme categories cannot be easily surmised from this visualization.

To estimate the relative distribution of gene detections, hypothesized to be bimodal given the histogram above, we compute the probability density function of the observed detection. This process takes the observed distribution and estimates a function describing the probability of observing any given value, such that the integral of any bounded x range equals the probability of observing that x in the observed data. The y-axis for density plots is scaled such that the total integral of the function equals one.

First, to prove that this bimodal tendency of most genes being either completely undetected (and therefore unambiguously EAG) or mostly detected, we plot the distibution of gene detection without separating ECG and EAG genes. The probability density function is produced and plotted with the following code (note that the bandwidth is set to 0.01 to allow for the sharp dropoff observed from the histogram above):

```
ggplot(many_ecag_genomes_long, aes(x = value, y = genome, height=..density..)) + 
  stat_density_ridges(color='black', geom="density_ridges_gradient", bandwidth = 0.01, scale=7) + 
  scale_x_continuous(expand = c(0,0)) + scale_y_discrete(expand = c(0,0)) +
  labs(x = "Detection (fraction of gene with > 0X coverage)", y = "Genome") +
  theme_classic()
```

This plot shows the probability (height) of observing a given gene detection (x axis) for all genes in each genome. Each different row is a different genome’s genes, and the height of the curve for a given detection value corresponds to the probability of observing genes with that detection value, based on all detection values obtained for all metagenomes for all genes with that detection.

Thus, we can be confident that genes most frequently to fall into two categories - mostly detected in many metagenomes, or completely undetected in many metagenomes. So, the large majority of genes we hope to identify as being environmentally accessory are absent from the majority of metagenomes and thus will always be identified as accessory regardless of any non-zero threshold used.

We then split the gene detections into two groups, based on whether they were ECG or EAG:

```
ggplot(many_ecag_genomes_long, aes(x = value, y = genome, height=..density..)) + 
  stat_density_ridges(aes(fill=ECAG), geom="density_ridges_gradient", bandwidth = 0.01, scale=7, lwd=0.2) +  
  scale_x_continuous(expand = c(0,0)) + scale_y_discrete(expand = c(0,0)) +
  labs(x = "Detection (fraction of gene with > 0X coverage)", y = "Genome", fill="EAG or ECG") +
  scale_fill_manual(values = c("#Fb4a2a7a", "#00AaFf7a")) +
  theme_classic()
```

From this plot, the detection remains clearly bimodal (most genes are either completely detected or completely undetected). The majority of genes are either ECG and fully or near-fully detected in the majority of metagenomes, or else EAG and completely absent (detection of zero) or nearly so from the majority of metagenomes. Thus, the specific value of the threshold between ECG and EAG is not critical, as the distribution is so sharply bimodal that picking any point between either extreme of the distributionhas little impact.

##### Combining metapangenomes

We then combined the metagenomes’ environmental core/accessory layer by habitat onto a single pangenome figure so we could directly compare the environmental representation of each homologous gene across sites.

All of each pangenome’s miscellaneous data were exported using `anvi-export-misc-data` parameter, e.g.

```
#!/bin/bash

prefix="Haemophilus-isolates"
readsStart=11  # which column in anvio layers table is the start of the metagenomes

sites=(td bm supp)  # important that td is first if that will be the one onto which others are added

for site in ${sites[*]}; do

anvi-export-misc-data -p $prefix-$site-PAN/$prefix-$site-PAN.db -t items -o $prefix-$site-items.txt
anvi-export-misc-data -p $prefix-$site-PAN/$prefix-$site-PAN.db -t layers -o $prefix-$site-layers.txt

# delete the unprocessed columns from TD so don't end up with two sets of TD metagenomes (formatted + unformatted)
anvi-delete-misc-data -p $prefix-td-PAN/$prefix-td-PAN.db -t layers --keys-to-remove $(head -1 $prefix-$site-layers.txt | cut -f$readsStart- - | tr '\t' ',')

# select metapan rings from items data and add oral habitat as prefix
awk -F"\t" -v site=$site 'BEGIN{site=toupper(site)}; NR==1{print $1 FS site"-"$9 FS site"-"$10 FS site"-"$11 FS site"-"$12} NR>1 {print $1 FS $9 FS $10 FS $11 FS $12}' $prefix-$site-items.txt > $prefix-$site-items.tmp
# prepend capitalized site ID before each metagenome's coverage of the genomes
cut -d' ' -f1,$readsStart- $prefix-$site-layers.txt | sed "s/SRS/\U$site-SRS/g" > $prefix-$site-layers.tmp

# trim off junk from SRS filenames in header
sed -i"" 's/_DENOVO_DUPLICATES_MARKED_TRIMMED//g' $prefix-$site-layers.tmp

# import this data back into the td data
anvi-import-misc-data -p $prefix-td-PAN/$prefix-td-PAN.db -t items $prefix-$site-items.tmp
anvi-import-misc-data -p $prefix-td-PAN/$prefix-td-PAN.db -t layers $prefix-$site-layers.tmp

done
```

For the `*layers.txt` file, which contains each genome’s per-sample coverage information, we used a custom R script to calculate and export the median coverage for each genome for each site, as well as combine the per-sample coverages for a heatmap:

We chose median as the most relevant measure of central tendency, since the presence of one very deeply sampled metagenome, or one metagenome with sampling contamination (e.g., a sample of plaque included saliva) could skew the mean.

```
t <- read.csv("Haemophilus-isolates-td-layers.tmp", sep = "\t", header = T)
b <- read.csv("Haemophilus-isolates-bm-layers.tmp", sep = "\t", header = T)
p <- read.csv("Haemophilus-isolates-supp-layers.tmp", sep = "\t", header = T)

# save genome ids for later
genomeIDs <- t[,1]

# get just the coverages
t <- t[,grep("SR", colnames(t))]
b <- b[,grep("SR", colnames(b))]
p <- p[,grep("SR", colnames(p))]

# get the median for each row=genome
t <- apply(t, 1, median)
b <- apply(b, 1, median)
p <- apply(p, 1, median)

# save to import into anvio
write.table(cbind(layers=as.character(genomeIDs), TD_HMP_MEDIAN=t, BM_HMP_MEDIAN=b, SUPP_HMP_MEDIAN=p), "haem_hmp_median.tsv", sep = "\t", col.names = T, row.names = F, quote=F)
```

which makes a file that looks like:

```
##      layers                                 TD_HMP_MEDIAN     
## [1,] "Haemophilus_parainfluenzae_1128_HPAR" "3.50155084141615"
## [2,] "Haemophilus_parainfluenzae_1209_HPAR" "6.815398696077"  
## [3,] "Haemophilus_parainfluenzae_137_HINF"  "3.1613550551289" 
## [4,] "Haemophilus_parainfluenzae_146_HPAR"  "15.1593914157683"
## [5,] "Haemophilus_parainfluenzae_155_HPAR"  "3.47668191817337"
## [6,] "Haemophilus_parainfluenzae_174_HPAR"  "3.35610746823065"
##      BM_HMP_MEDIAN       SUPP_HMP_MEDIAN    
## [1,] "0.657201515386123" "1.06875757236494" 
## [2,] "1.08419195652694"  "1.55167011239881" 
## [3,] "0.565412049050462" "1.9723758712279"  
## [4,] "1.06501435940931"  "1.88690992300501" 
## [5,] "0.563448807228174" "0.794885888409524"
## [6,] "0.636198880422302" "2.31254813948498"
```

All these data were then imported into the TD pangenome database (TD chosen arbitrarily) using `anvi-import-misc-data -t layers`.

##### Anvi’o interactive display choices for metapangenomes

The genome layers in the pangenomes were ordered by gene cluster frequency, and the gene clusters were ordered by frequency in genomes.

Coloring and spacing were set manually through the anvi’o interactive interface `anvi-display-pan`. Spacing between genomes in the pangenome was done to improve the eye’s ability to follow groups identified by the gene cluster frequency dendrogram (setting the layer order) throughout the pangenome.

##### Supplemental tables

Supplemental tables of gene clusters and their functions (i.e., SD1, SD3 in the paper, which are the contents of pangenomes in tabular format) were generated by summarizing each combined metapangenome with `anvi-summarize`, like

```
anvi-summarize -g Haemophilus-isolates-GENOMES.db -p Haemophilus-isolates-td-PAN/*PAN.db -C default -o Haemophilus-isolates-td-PAN-SUMMARY
```

#### Querying the gene cluster annotations for enriched functions

Pangenomes ultimately rely on the ability to cluster genes into ‘gene clusters’ based on detecting islands in amino-acid sequence space (described earlier in “Pangenome creation methods details”). For the conceptual and evolutionary logic of what gene clusters represent, please see the Supplemental Text of our manuscript. But, the biological interpretation of a gene cluster requires investigation. That is, we wish to know whether gene clusters generally correspond to represent distinct functions, or whether genes of the same function are split into multiple groups based on shared similarities. This has significant implications for how one interprets the core genome vs. accessory genome.

The fundamental challenge of creating a pangenome is that each gene may be under a different selective regime. For instance, a ribosomal protein evolves under different constraints than a given transcriptional regulator or than a membrane transporter. An extreme example is the gene encoding the RuBisCO large subunit, all of which fix CO\_2\_ yet the sequences are divergent except for a tiny window of conservation around the active site. On the other hand, the 16S ribosomal RNA gene sequence is much more conserved. And, even a partial truncation in some genes can totally abolish enzymatic function (e.g. Sirias et al. 2020).

To understand the functional implications of our gene clusters, we took two approaches: 1) Re-create the pangenome, but plot functions on the radial axis vs genomes, instead of gene clusters x genomes (most useful for Rothia). 2) Count up the number of gene clusters per function, and pay special attention to cases where one function is found in all genomes but split into species-specific gene clusters.

To re-create the pangenome, we applied Alon Schaiber’s example code to our data. This process finds functions enriched in each of the defined groups. Here, we defined groups as the three subgroups of H. parainfluenzae detected in the pangenome. The core program of this step is `anvi-get-enriched-functions-per-pan-group` which identifies functions and their frequencies. This step also creates the functional enrichment tables, the specifics of which will be discussed in the next section.

Following the steps in Alon Schaiber’s guide, this pangenome for Rothia displaying functions was created.

As expected, many genes from the singleton accessory genome disappear as they lack functional annotation, and the relative representation of the core genome is much larger since core genes are typically more studied and better annotated. Yet, the overall pattern of the pangenome remains visually similar, with a large core genome, species-specific core genomes, and the greater similarity between *R. dentocariosa* and *R. aeria* than either share with *R. mucilaginosa*.

To count up the functions by gene clusters, to identify places where the same function was split across multiple gene clusters (particularly if that split matched species groups, e.g., if anvi’o made three DnaK gene clusters, one for each species’ DnaK), we branched off from this workflow. When we ran `anvi-summarize` on the pangenomes above, they produced files named like `Haemophilus-isolates-td-PAN-SUMMARY/Haemophilus-isolates-td_gene_clusters_summary.txt` that among other things report the annotation for each gene and which gene cluster to which that gene belongs. From this, we can count up how many times each function was found, by core or accessory set. Here is the Python we used to parse this file:

```
#!/usr/bin/env python3

import pandas as pd

summaryPath = 'Haemophilus-isolates-td-PAN-SUMMARY/Haemophilus-isolates-td_gene_clusters_summary.txt'

summary = pd.read_csv(summaryPath, sep="\t", dtype={'ProDom_ACC': str, 'ProDom': str})

summary = summary[['gene_cluster_id','bin_name','functional_homogeneity_index','geometric_homogeneity_index', 'TIGRFAM','Pfam']]

summary = summary[pd.notnull(summary['Pfam'])]


funcCounts = []

for pfam in summary['Pfam'].unique():
    subPfam = summary.ix[summary['Pfam'] == pfam]
    for bin in subPfam['bin_name'].unique():
        subPfamsubBin = subPfam.ix[subPfam['bin_name'] == bin]
        gcIDsUnique = list(subPfamsubBin['gene_cluster_id'].unique())
        df = pd.DataFrame({'bin_name': [bin], 'Pfam': [pfam], 'num_uniq_gcs': [len(gcIDsUnique)],
                                'uniq_gc_ids': [','.join(gcIDsUnique)]})
        funcCounts.append(df)

funcCounts = pd.concat(funcCounts, axis=0)
funcCounts.to_csv(summaryPath + "-homogeneity.tsv", sep="\t", index=None)
```

We now switch from *H. parainfluenzae* to the genus *Rothia* since having genus vs species is more illustrative for how gene clusters treat the varying levels of amino acid similarity within a species vs within a genus.

This produced a slimmed-down table with only the relevant information, but it was still difficult to conceptualize all these data. We plotted this information and generated a wide-format table providing a breakdown, for each function, of how many gene clusters existed, and the set of core genes to which they belonged (e.g., genus core, aeria + dentocariosa core, singleton accessory genome, etc.).

```
# plot pfam redundancy of GCs for rothia metapangenome

library(ggplot2)
library(reshape2)

funcCounts <- read.csv('Rothia-isolates-HMP_gene_clusters_summary.txt-homogeneity.tsv', sep = "\t")

funcCounts <- funcCounts[,!(colnames(funcCounts) %in% "X")]

coreCols <- setNames(c('green','blue','turquoise','black','pink','purple','orchid','magenta','grey80','grey50'),levels(funcCounts$bin_name))

funcCounts$Pfam <- factor(funcCounts$Pfam, levels = unique(as.character(funcCounts$Pfam[order(funcCounts$bin_name)])))  # make cores clump
funcCounts$Pfam <- factor(funcCounts$Pfam, levels = names(rev(sort(tapply(funcCounts$num_uniq_gcs, funcCounts$Pfam, sum)))))  # order by decreasing total GC count

ggplot(funcCounts, aes(x = Pfam, y = num_uniq_gcs)) +
    geom_bar(stat = 'identity', position = 'stack', aes(fill = bin_name), width = 1) +
    theme_minimal() + scale_x_discrete(expand = c(0,0)) + scale_y_continuous(expand = c(0,0)) +
    scale_fill_manual(values = coreCols) + labs(y = "Num. gene clusters", fill = 'Bin') +
    theme(panel.grid = element_blank(), panel.background = element_rect(fill = NA, color = 'black'), axis.text = element_text(color = 'black'),
          axis.text.x = element_text(angle = 90, hjust = 1, size = 1))
```

Here are the first few rows of the table:

```
funcTable <- dcast(funcCounts, Pfam ~ bin_name, value.var = 'num_uniq_gcs')
funcTable[is.na(funcTable)] <- 0

head(funcTable, 30)
```

```
##                                                                   Pfam a_452
## 1                                                      ABC transporter     3
## 2                                      Pentapeptide repeats (9 copies)     5
## 3                                        Major Facilitator Superfamily     1
## 4                                                           AAA domain     3
## 5                                              Helix-turn-helix domain     1
## 6                           Bacterial regulatory proteins, tetR family     1
## 7                                              S-layer homology domain     4
## 8               Type I restriction modification DNA specificity domain     0
## 9                                                           RHS Repeat     0
## 10                                                   N-6 DNA Methylase     0
## 11                                       Glycosyl transferase family 2     1
## 12                                                        AIPR protein     0
## 13                            Type III restriction enzyme, res subunit     0
## 14                                          Zinc-binding dehydrogenase     0
## 15                                                        NUDIX domain     0
## 16                                     Acetyltransferase (GNAT) family     2
## 17                                                   AAA ATPase domain     0
## 18                                                       DNA methylase     0
## 19                                                  AMP-binding enzyme     0
## 20                                            Methyltransferase domain     0
## 21                                    Excalibur calcium-binding domain     4
## 22                                       Glycosyl transferases group 1     2
## 23                                              Acyltransferase family     2
## 24 Binding-protein-dependent transport system inner membrane component     0
## 25                                                    Helix-turn-helix     0
## 26                                           alpha/beta hydrolase fold     0
## 27                                                      Fic/DOC family     1
## 28                                                         MarR family     1
## 29                                  Protein of unknown function DUF262     0
## 30                                                    HNH endonuclease     0
##    d_267 da_204 g_1130 m1_39 m2_38 m3_22 mAll_205 s_1371 UNBINNED_ITEMS_BIN
## 1      0      3     27     0     0     0        3      0                 28
## 2      0      0      0     0     0     0        0      9                 36
## 3      0      2      7     0     0     0        1      2                 21
## 4      2      0      7     0     0     0        1      7                  9
## 5      0      1      4     0     0     0        1      3                 19
## 6      2      4      6     0     0     0        1      2                 11
## 7      4      0      2     1     0     0        5      0                 11
## 8      0      0      0     0     0     1        0     12                 13
## 9      0      0      0     0     0     0        0      5                 18
## 10     0      0      0     0     0     0        0      3                 18
## 11     1      1      8     0     0     0        0      0                  8
## 12     0      0      0     0     0     0        0      8                  6
## 13     0      0      2     0     0     0        0      2                  9
## 14     0      3      6     0     0     0        0      0                  4
## 15     0      1     10     0     0     0        0      0                  2
## 16     1      3      3     0     0     0        0      0                  4
## 17     0      0      0     0     0     0        0      4                  8
## 18     0      0      1     0     0     0        0      3                  8
## 19     0      0      8     0     0     0        0      3                  1
## 20     0      2      3     0     0     0        0      2                  5
## 21     2      0      0     0     0     0        2      0                  4
## 22     0      0      8     0     0     0        0      0                  2
## 23     0      1      2     0     0     0        0      0                  7
## 24     0      0      5     0     0     0        1      0                  5
## 25     0      1      0     0     0     0        0      3                  7
## 26     1      1      2     0     0     0        0      1                  6
## 27     0      0      0     0     0     0        0      0                 10
## 28     0      3      1     0     0     0        0      2                  4
## 29     0      0      0     0     0     0        0      6                  4
## 30     0      0      0     0     0     0        0      3                  7
```

#### Differential function analysis

To investigate potential functional drivers affecting the differential abundances across habitats evidenced by *H. parainfluenzae* strains, we assigned each genome to one of the three genomic groups observed from the metapangenome (Figure 2 in text) and looked for functions enriched in a particular group relative to the other groups.

Here is the file showing each genome’s assignments:

```
layer   habitat
Haemophilus_parainfluenzae_C2004002727  Group3
Haemophilus_parainfluenzae_ATCC_9796    Group3
Haemophilus_parainfluenzae_C2005004058  Group3
Haemophilus_parainfluenzae_UMB0748      Group3
Haemophilus_parainfluenzae_C2006002596  Group3
Haemophilus_parainfluenzae_CCUG_58848   Group3
Haemophilus_parainfluenzae_488_HPAR     Group3
Haemophilus_parainfluenzae_ATCC_33392_v1        Group3
Haemophilus_parainfluenzae_ATCC_33392_v2        Group3
Haemophilus_parainfluenzae_174_HPAR     Group3
Haemophilus_parainfluenzae_137_HINF     Group3
Haemophilus_parainfluenzae_HK2019_HK2019        Group3
Haemophilus_parainfluenzae_HK262_HK262  Group3
Haemophilus_parainfluenzae_CCUG_62654   Group3
Haemophilus_parainfluenzae_CCUG_62655   Group3
Haemophilus_parainfluenzae_C2004000280  Group2
Haemophilus_parainfluenzae_C2004002729  Group2
Haemophilus_parainfluenzae_901_HPAR     Group2
Haemophilus_parainfluenzae_432_HPAR     Group2
Haemophilus_parainfluenzae_146_HPAR     Group2
Haemophilus_parainfluenzae_209_HPAR     Group2
Haemophilus_parainfluenzae_T3T1_T3T1    Group2
Haemophilus_parainfluenzae_215035_2_ISO5        Group2
Haemophilus_parainfluenzae_C2008001710  Group2
Haemophilus_parainfluenzae_C2008003258  Group1
Haemophilus_parainfluenzae_C2009038101  Group1
Haemophilus_parainfluenzae_C2011020591  Group1
Haemophilus_parainfluenzae_60884_B_Hi_2 Group1
Haemophilus_parainfluenzae_65114_B_Hi_3 Group1
Haemophilus_parainfluenzae_777_HPAR     Group1
Haemophilus_parainfluenzae_155_HPAR     Group1
Haemophilus_parainfluenzae_1128_HPAR    Group1
Haemophilus_parainfluenzae_1209_HPAR    Group1
```

We called this `habitat_groups.txt` and then loaded it into the anvi’o pangenomes DB with

```
anvi-import-misc-data -p Haemophilus-isolates-td-PAN/Haemophilus-isolates-td-PAN.db -t layers habitat_groups.txt
```

We then used the `anvi-get-enriched-functions-per-pan-group` command for both Pfam and TIGRfam annotations in the following way:

```
anvi-get-enriched-functions-per-pan-group -p Haemophilus-isolates-td-PAN/Haemophilus-isolates-td-PAN.db -g Haemophilus-isolates-td-GENOMES.db --category habitat --annotation-source TIGRFAM -o Haemophilus-isolates-enriched-functions-habitat-tigr.txt
```

This produces Supplemental Data 2 (TIGRFAM). Note that this snippet is specifically for TIGRFAMs; we ran it also again with `--annotation-source Pfam` to get a table of differential Pfam functions, and similarly for the *Rothia* pangenome with COG functional categories to identify major functional differences and similarities between species.

#### Visualizing mapping results at the per-gene, per-sample level

To get a better idea of what the per-gene coverages looked like along with the metapangenomes, we extracted the per-gene coverage information for each genome.

In contrast to the summary metric of coverage that reports a genome-wide mean, this step permits visualizing the distribution of coverage within a genome to estimate the quality of coverage. This approach allows the identification of human contaminant genes, as such genes would receive much more coverage than the rest of the genome and thereby bias the overall coverage obtained byt he genome. Also, the evenness of coverage, and the max/min ratio of coverage at the gene level informs many of the previous choices (such as whether the specific threshold value matters for defining environmental core vs. environmental accessory genes, leading to the gene-level detection described above).

Biologically, inspecting the per-gene, per-sample coverage within a genome offers a means to estimate how similar the population in the metagenomes are to the reference. If the metagenome contains multiple closely related populations that each containing a non-overlapping subset of genes shared with a given cultivar genome, the genome’s detection (fraction of genes or nucleotides covered) will be high, but each gene’s coverage might be very different. On the other hand, if the metagenome only contains a single dominant population related to the reference, then most of the genome will receive coverage, and the variance of the coverage from gene to gene should be lower than in the first scenario. Alternatively, if the metagenome does not contain a population similar to the genome of interest, then most of the genes in the genome will not receive coverage (since there are no populations contributing those genes to the metagenome), but mobile elements or a few highly conserved genes such as rRNAs or ribosomal proteins may receive coverage reads originating from distantly related bacteria. In this last scenario the genome will have low detection and low coverage, and plots of each gene’s coverage would look mostly empty with the occasional spike of coverage.

Importantly, this per-gene, per-sample information is independent of the system and so is applicable for microbial populations outside the human mouth.

This per-gene, per-sample coverage is shown in Figure 4A, which was created with the following code (so using the Rothia dataset for this example code). This was done in the background in `anvi-meta-pan-genome` and can be done ad-hoc with `anvi-interactive --gene-mode`, but we needed more control for formatting and miscellaneous data so we re-gathered the nucleotide-level coverage information per genome, and stored this information individually. We used the following script to collect and organize the data:

```
#!/bin/bash
#SBATCH -N 1 #1 nodes of ram
#SBATCH -n 1 # 1 cores from each
#SBATCH --contiguous
#SBATCH --mem=24G #per node
#SBATCH -t 0-5:00:00
#SBATCH -p shared
#SBATCH --array=0-68%69
#SBATCH --job-name="MP-$target"
#SBATCH -o odyssey_meta-$target.out
#SBATCH -e odyssey_meta-$target.err
#SBATCH --mail-type=NONE

taxPrefix="Rothia-isolates"

### RUN THESE 2 LINES FIRST
#mkdir genomesMapped
#awk '{print $2}' $taxPrefix-renaming-manual.tsv | uniq | split -l 1 -d -a 3 - genomesMapped/genome-

cd genomesMapped

taskNum=$(printf %03d $SLURM_ARRAY_TASK_ID)

# which genome this batch is targeting
target=$(cat genome-$taskNum)

# activate anvio
source ~/virtual-envs/anvio-dev-venv/bin/activate

## for each habitat, make a dir, enter, get this genome's coverage and detection info, then go back up one level to do them all
mkdir $target-TD
cd $target-TD
anvi-script-gen-distribution-of-genes-in-a-bin -c ../../$taxPrefix-CONTIGS.db -p ../../$taxPrefix-td-MERGED/PROFILE.db -C Genomes -b $target --fraction-of-median-coverage 0.25
cd ..

mkdir $target-PQ
cd $target-PQ
anvi-script-gen-distribution-of-genes-in-a-bin -c ../../$taxPrefix-CONTIGS.db -p ../../$taxPrefix-supp-MERGED/PROFILE.db -C Genomes -b $target --fraction-of-median-coverage 0.25
cd ..

mkdir $target-BM
cd $target-BM
anvi-script-gen-distribution-of-genes-in-a-bin -c ../../$taxPrefix-CONTIGS.db -p ../../$taxPrefix-bm-MERGED/PROFILE.db -C Genomes -b $target --fraction-of-median-coverage 0.25
cd ..
```

This script was run in the top level directory, where all the other files were generated. Note that this is an array, so the two commented out lines (`mkdir` and `awk`) were run first as one-liners to set everything up. This script then creates a set of batch information inside that `genomesMapped` directory, which will also have 3 directories for each genome, one for each habitat. The `--fraction-of-median-coverage` parameter was kept at `0.25`, identical to the metapangenome calculation above.

While the main text of the manuscript describes this analysis for only *R.* sp. E04 and *R.* sp. C03, we generated and inspected such per-gene, per-sample visualizationss for each genome.

This output of this script made separate directories for each genome for each site. Ideally, we would visualize data from all the metagenomes, but in this case with HMP data, 551 metagenomes are far too many to distinguish on the average size screen. So, we select the top N samples (top being determined by highest median coverage) from each site for each genome and combine into a single figure, along with the environmental core/accessory designations for each gene from each habitat. We wrote a helper python script to help with the merging:

```
#!/usr/bin/env python

from optparse import OptionParser
import pandas as pd
import os

parser = OptionParser()
(options, args) = parser.parse_args()

fileRoot=args[0]
keep=int(args[1])

# load coverages
covs_td = pd.read_csv("../"+args[0]+"-TD/"+args[0]+"-GENE-COVs.txt", sep ="\t", index_col=False)
covs_bm = pd.read_csv("../"+args[0]+"-BM/"+args[0]+"-GENE-COVs.txt", sep ="\t", index_col=False)
covs_supp = pd.read_csv("../"+args[0]+"-PQ/"+args[0]+"-GENE-COVs.txt", sep ="\t", index_col=False)

# load detections
det_td = pd.read_csv("../"+args[0]+"-TD/"+args[0]+"-ENV-DETECTION.txt", sep ="\t", index_col=False)
det_bm = pd.read_csv("../"+args[0]+"-BM/"+args[0]+"-ENV-DETECTION.txt", sep ="\t", index_col=False)
det_supp = pd.read_csv("../"+args[0]+"-PQ/"+args[0]+"-ENV-DETECTION.txt", sep ="\t", index_col=False)

det_td.columns.values[1] = 'TD_detection'
det_bm.columns.values[1] = 'BM_detection'
det_supp.columns.values[1] = 'SUPP_detection'

# sort by max median coverage
covs_td = covs_td.reindex_axis(covs_td.median().sort_values(ascending=False).index, axis=1)
covs_bm = covs_bm.reindex_axis(covs_bm.median().sort_values(ascending=False).index, axis=1)
covs_supp = covs_supp.reindex_axis(covs_supp.median().sort_values(ascending=False).index, axis=1)

# add in sites to samples
covs_td.columns.values[1:] = [x+"-TD" for x in covs_td.columns.values[1:]]
covs_bm.columns.values[1:] = [x+"-BM" for x in covs_bm.columns.values[1:]]
covs_supp.columns.values[1:] = [x+"-SUPP" for x in covs_supp.columns.values[1:]]

# make the new coverages
covs_keep = pd.concat([covs_td.iloc[:,0:(keep+1)], covs_bm.iloc[:,1:(keep+1)], covs_supp.iloc[:,1:(keep+1)]], axis=1)
det_all = pd.concat([det_td, det_bm.iloc[:,1], det_supp.iloc[:,1]], axis=1)

# save it
covs_keep.to_csv(args[0]+"-GENE-COVs.txt-COMBO-"+str(keep), sep="\t", index=False)
det_all.to_csv(args[0]+"-ENV-DETECTION.txt-COMBO-"+str(keep), sep="\t", index=False)

keys = sorted(covs_keep['key'].tolist(), reverse=True)
with open("synteny.txt", 'w') as f:
        for k in keys:
                f.write(str(k)+'\n')


with open("RUN_ME_FOR_ANVIO", "w") as f:
        f.write("anvi-interactive -P 8081 --manual --title '{0}' -d {0}-GENE-COVs.txt-COMBO-{1} -A {0}-ENV-DETECTION.txt-COMBO-{1} -p {0}-PROFILE.db --items-order synteny.txt".format(args[0],keep))

os.chmod("RUN_ME_FOR_ANVIO", 0o755)
```

We named this script `indivMetaCombiner.py` in each directory and ran it with `./indivMetaCombiner.py Rothia_sp_HMSC061E04_HMSC061E04 30`, where the first argument is the file prefix for the earlier environmental detection output, which also happens to be the name of the genome, and the second argument is the number of samples to include (N), taking the N with the highest median coverage across samples for that habitat. We chose median, since mean could be biased by one outlier gene (e.g. 16S) picking up reads, and we are not interested displaying any samples governed by such an outlier.

#### Nucleotide level coverage plots (Figure 4A, 4B)

As for the gene-level coverage above, the nucleotide-level coverage and variability over genes of interest is useful for determining whether the coverage comes from a close relative, a distant relative, or is potentially an artifact. The amount of nucleotide variants mapped also informs this inference. For example, if nucleotide-level coverage is relatively even, with few single nucleotide variants (SNVs), then the reads likely come from a closely-related population. If the coverage is relatively even yet most of the mapped reads contain SNVs, then the population providing those reads likely is more distantly related to the reference gene. Finally, if the coverage of a gene has many gaps, representing subregions in the gene receiving no coverage, and the regions recruiting coverage do so with many SNVs, then likely the coverage results from distantly related conserved domains or chance similarity and does not signify the presence of that gene in a closely-related population.

###### Rothia candidate drivers of adaptation to BM

After identifying a few candidate drivers of habitat specificity for *R. mucilaginosa* populations living in buccal mucosa, we used the interactive anvi’o interface `anvi-interactive` with `--gene-mode` to identify the contig and split from which the gene came.

For each oral habitat, we exported the coverage using

```
anvi-get-split-coverages -p $prefix-$site-MERGED/PROFILE.db -C Genomes -b Rothia_sp_HMSC061E04_HMSC061E04 -o rot_split_cov-ALL-$site.tsv
```

changing the path as needed for each habitat’s data.

Then, similarly adjusting the path to specify each habitat, we obtained the sequence variants mapped from the metagenome for each nucleotide position for the contig.

```
anvi-gen-variability-profile -c $prefix-CONTIGS.db -p $prefix-$site-MERGED/PROFILE.db -C Genomes -b Rothia_sp_HMSC061E04_HMSC061E04 -o rot_split_cov-ALL-variability-$site.tsv
```

To link genes to split-based coverage, we exported the table listing the split-based start/stop positions for each gene with sqlite as follows:

```
sqlite3 Haemophilus-isolates-CONTIGS.db 'PRAGMA table_info(genes_in_splits)' | awk -F"|" '{print $2}' | tr '\n' '|' | sed "s/|$//" > Rothia-isolates-splits-genes.txt
sqlite3 Rothia-isolates-CONTIGS.db 'SELECT * from genes_in_splits' >> Rothia-isolates-splits-genes.txt
```

With these inputs, we can plot per-nucleotide coverage from the metagenomes on a window around a focal gene, as in Figure 4b.

```
library(ggplot2)
library(reshape2)
library(entropy)
library(dplyr)

# read in and pre-process variability information
getVariability <- function(prefix, sites=c('TD','BM','SUPP'), pos.min = 0, pos.max = 999999, min_coverage=0) {
    all <- data.frame()

    # iteratively load and attach row-wise
    for (site in sites) {
        df <- read.csv(paste0(prefix, "variability-", site, '.tsv'), sep = "\t", header = T)
        
        # only keep variability info for sites with > 10x coverage
        df <- df[df$coverage >= min_coverage,]
        
        # get entropy across samples and lose junk
        df <- group_by(df, split_name, pos) %>% summarise(a=sum(A), c=sum(C), t=sum(T), g=sum(G)) %>%  # collapse so have split, position in split, and sum of samples
          group_by(split_name,pos) %>% summarise(entropy=entropy.empirical(c(a,c,t,g), unit = 'log2'))  # regroup to keep split & pos, then entropy across all samples
        
        df <- data.frame(df)
        
        df$site <- toupper(site)

        all <- rbind(all, df)
    }

    # subset if requested
    all <- all[(all$pos >= pos.min) & (all$pos <= pos.max),]

    all$site <- factor(all$site, levels = c("SUPP","BM","TD"))

    return(all)
}


getCoverages <- function(prefix, sites=c('TD','BM','SUPP'), pos.min = 0, pos.max = 999999) {
    all <- data.frame()

    # iteratively load and attach row-wise
    for (site in sites) {
        df <- read.csv(paste0(prefix, site, '.tsv'), sep = "\t", header = T)#, nrows = 1000000)
        colnames(df) <- c("unique_pos_id","nt_position","split_id","sample_name","coverage")
        
        df$site <- toupper(site)

        all <- rbind(all, df)
    }

    return(all)
}

# THIS FIGURES OUT WHICH SPLITS NEED COVERAGE - doing this first to reduce filesize of the covs file(s)
findSpecificSplits <- function(geneID="104649", genes) {
  
  # find what contig the gene is on
  splitName <- genes$split[genes$gene_callers_id == geneID]
  splitNum <- unlist(strsplit(as.character(splitName), "_"))[4]
  splitBody <- unlist(strsplit(as.character(splitName), "_"))[1:3]
  
  # get adjacent splits too just in case it's on edge of split (same contig, though)
  adjacentSplits <- sprintf(paste0(c(splitBody, "%05s"), collapse = "_"), (as.numeric(splitNum)-1):(as.numeric(splitNum) + 1))
    
  
  splits <- unique(as.character(genes$split[genes$split %in% adjacentSplits]))
  
  return(splits)
}

plotCovsAndGenes <- function(covs = getCoverages(), genes = getGeneCalls(), surroundingGenes=10, prefix="", variability = NULL,
                             tax, splits, keep_fraction = 1, target, site="TD", outputSuf=NULL) {
  
  splitID <- genes$split[genes$gene_callers_id == target]
    
  workingCov <- covs[grep(splitID, covs$split_id),]

  # take n genes either side
  workingGcs <- genes[(which(genes$gene_callers_id == target) - surroundingGenes):(which(genes$gene_callers_id == target) + surroundingGenes),]

  # make sure they're all on same contig
  workingGcs <- workingGcs[workingGcs$split == workingGcs$split[workingGcs$gene_callers_id == target],]

  # pull out window provided by GeneCaller positions
  workingCov <- workingCov[(workingCov$nt_position >= workingGcs$start_in_split[1]) & 
                             (workingCov$nt_position <= workingGcs$stop_in_split[length(workingGcs$stop_in_split)]),]
  
  print(head(workingCov))
  
  genes <- workingGcs
  covs <- workingCov
  
    genes$site <- factor(site)
    genes$gene_callers_id <- factor(genes$gene_callers_id)

    geneColors <- setNames(rep("grey80", nrow(genes)), as.character(genes$gene_callers_id))
    geneColors[target] <- "#BA0000"
    
    ###############
    # drop variability info not in window, fix NAs to 0 for scaling
    variability <- variability[variability$split_name %in% unique(as.character(covs$split_id)),]
    
    # sometimes chunk might not have variability and crash; only renormalize if kept any
    if (nrow(variability) > 0){
      print("found variability; proceeding...")
      variability <- variability[(variability$pos >= min(covs$nt_position)) & (variability$pos <= max(covs$nt_position)),]
      variability$site <- factor(variability$site, levels = c("SUPP","BM","TD"))
      variability$entropy[is.na(variability$entropy)] <- 0
      
      # multiply out coverage to get proportional SNPs
      variability$entropyOrig <- variability$entropy
      variability$entropy <- variability$entropy * (max(covs$coverage)/max(variability$entropy))
      
      print(max(variability$entropyOrig))
    } else { # make a dummy
      print("no variability in this window; making a dummy")
      variability[1, ] <- NA
      variability$site <- site
      variability$entropyOrig <- 0
    }


  #######
    # make the plot
    p <- ggplot(data = covs, aes(x = nt_position, y = coverage)) +
        facet_wrap(~site, ncol = 1, scales = 'fixed', strip.position = 'right', ) + theme_minimal() +
        geom_col(data = variability, stat = 'identity', position = 'identity', aes(x = pos, y = entropy), color = 'black', alpha = 0.5, size = 0.2) +
        geom_line(size = 0.3, alpha = 0.5, aes(color=site, group=sample_name)) +
        geom_hline(data = genes, yintercept = 0.97*max(covs$coverage), color = 'grey20') +
        geom_segment(data = genes, aes(x = start_in_split, xend = stop_in_split, color = gene_callers_id), 
                     y = 0.99*max(covs$coverage), yend = 0.99*max(covs$coverage), size = 4) +
        labs(title = target) + ylab("Coverage") +
        scale_x_continuous(expand=c(0,0)) + scale_y_continuous(expand=c(0,0), sec.axis = sec_axis(~.*max(variability$entropyOrig)/max(covs$coverage))) +
        scale_color_manual(values = c(setNames(c("#2790db", "#b287e8", "#69d173"), c("TD","BM","SUPP")), geneColors)) + guides(color = F) +
        theme(panel.grid = element_blank(), axis.line = element_line(colour = 'black', size = 0.5), axis.text = element_text(colour='black'),
              strip.text = element_text(angle=90), axis.ticks = element_line(colour='black', size=0.3))


    # did you give me a suffix to use to save it?
    if (!is.null(outputSuf)){
        pdf(paste0(prefix,outputSuf,".pdf"), width = 7, height = 4)
    }

    print(p)

    if (!is.null(outputSuf)){
        dev.off()
    }
}

prefix <- '~/g3/oral_genomes_analysis/rothia/rot_split_cov-ALL-'

geneCallsPath <- 'Rothia-isolates-splits-genes.txt'

tax <- 'R. muciaginosa E04'

splitPosRelativeToContig <- 203284 # rothia_buccal

geneTarget <- c("104283","103512","104082","103409","103186","103758","103125","104483","104321","104823","104402","103432",
                "104322","104458","104363","103468","103939","103201","104081","104464","104323","103386","104649","103463")  # 24 e04 genes from 22 gcs


# get gene call positions
gcs <- read.csv(geneCallsPath, header = T, sep="|")

# then find out what splits needed
splitsNeeded <- c()
for (gene in geneTarget) { 
  splitsNeeded <- c(splitsNeeded, findSpecificSplits(geneID = gene, genes = gcs))
}

site='SUPP'

# get coverages
cov <- getCoverages(prefix = prefix, pos.min = 0, pos.max = Inf, sites = site)
# and variability
variability <- getVariability(prefix = prefix, sites = site, min_coverage = 0)


# save each one
for (i in 1:length(geneTarget)) {
  plotCovsAndGenes(covs = cov, prefix=paste0('~/g3/oral_genomes_analysis/rothia/bm_gene_covs/20190826/E04_genes_',site, "-"),
                 genes = gcs, target = geneTarget[i],
                 variability = variability,
                 tax = tax, splits=splitDF, site = site,
                 outputSuf = geneTarget[i])
}
```

Which created the components of this figure.

These three panels show the gene highlighted in Figure 4B with 20 Kbp of context. Notice how TD coverage (bottom, each blue trace shows coverage from a different metagenome) is extremely low over the focal gene (rectangle highlighted in red), relative to the context, and where there are spikes of coverage allowing enough depth to estimate SNVs there are many SNVs. For the highlighted gene in particular, coverage is low and when there is a bump in coverage from a single metagenome, it covers only a fraction of the gene. This pattern cannot support the existence of this gene in TD populations.

On the other hand, the BM metagenomes (middle plot, purple traces) have consistently higher coverage across this window, including over the gene of interest. Many samples have high, even coverage over the highlighted gene, although not all. Also, where there is high coverage (over 100x in some metagenomes) there are few to no SNVs, indicating that these exact sequences were found in BM metagenomes abundantly, while that was not the case for TD. SUPP mapping is generally low except for one outlier metagenome with consistent high coverage. We interpret this pattern to indicate that E04 does not represent any abundant population in SUPP; the one sample contributing coverage may have resulted from from contamination during sampling, perhaps from the sampling instrument contacting the cheek during sampling of plaque. Regardless, these data show that this 20 Kbp context, and focal gene in particular, are abundant in *R. mucilaginosa* inhabiting the buccal mucosa, while rare in *R. mucilaginosa* inhabiting the tongue population.
